# A Human Alveolus-on-Chip Recapitulates SARS-CoV-2-mediated Lung Injury in an Organ-relevant Context for Pre-clinical Applications

**DOI:** 10.1101/2025.10.17.683096

**Authors:** Mirjam Kiener, Lea Lara De Maddalena, Nuria Roldan, Panagiotis Chouvardas, Laurène Froment, Marta De Menna, Thomas Geiser, Janick Stucki, Nina Hobi, Marianna Kruithof-de Julio

**Affiliations:** Urology Research Laboratory, Department for BioMedical Research (DBMR), University of Bern, Bern, Switzerland; Alveolix AG, Swiss Organs-on-Chip Innovation, Bern, Switzerland; Department of Urology, Inselspital, Bern University Hospital, University of Bern, Bern, Switzerland; Translational Organoid Resource (TOR), Department for BioMedical Research (DBMR), University of Bern, Bern, Switzerland; Department of Pulmonary Medicine, Allergology and Clinical Immunology, Inselspital, Bern University Hospital, University of Bern, Bern, Switzerland; Lung Precision Medicine, Department for BioMedical Research (DBMR), University of Bern, Bern, Switzerland; Organs-on-Chip Technologies, ARTORG Center for Biomedical Engineering, University of Bern, Bern, Switzerland

**Author notes:** Co-authorship (Nina Hobi, Marianna Kruithof-de Julio).

**Keywords:** Lung-on-Chip (LOC), COVID-19, SARS-CoV-2, alveolar barrier, pulmonary inflammation, drug efficacy testing

## Abstract

Respiratory viruses pose a constant threat to public health as highlighted by the pandemic outbreak of COVID-19. The most severe manifestation of COVID-19 is observed in the distal lung, where SARS-CoV-2 infection can result in massive inflammation and barrier breakdown. To date, few therapies are approved for clinical use in COVID-19 patients partially due to the lack of highly translational pre-clinical models of the alveoli. Human-derived microphysiological systems pose a promising new class of *in vitro* models to study viral infection in a relevant context. Therefore, we aimed to develop a Lung-on-Chip (LOC) model for studying SARS-CoV-2 infection at the alveolar barrier.

Using an immortalized alveolar epithelial cell line, ^AX^iAEC, and human lung microvascular endothelial cells (hLMVEC) we established a SARS-CoV-2 infection model on a LOC system. The LOC models the alveolar epithelial/endothelial barrier under physiological breathing motion.

Our results demonstrated that ^AX^iAEC maturation at the air-liquid interface (ALI) is essential for SARS-CoV-2 infection. SARS-CoV-2 infected the breathing LOC model and induced breakdown of the air-blood barrier. Finally, we evaluated the application of the LOC SARS-CoV-2 infection model for efficacy testing and demonstrated the antiviral effect of remdesivir. Drug treatment not only inhibited viral replication but also protected the alveolar barrier from damage and partially reverted SARS-CoV-2-mediated transcriptional dysregulation in ^AX^iAEC.

This novel LOC infection model recapitulates aspects of COVID-19. Our results highlight the importance of physiological cues such as ALI and stretch to accurately model host-pathogen interactions in the distal lung. Application of LOC models in pre-clinical drug testing may facilitate candidate compound selection at an early stage, allowing the allocation of resources to a few promising candidate compounds raising the potential to accelerate drug development and reduce costs and animal testing.

## Introduction

In late 2019, a mysterious series of viral pneumonia causing cough, fever, and dyspnoea emerged in Wuhan city, China, and was declared a pandemic outbreak of coronavirus disease 2019 (COVID-19) by the World Health Organization (WHO) in early 2020^1^. The causing agent of COVID-19, SARS-CoV-2, is a single-stranded positive-strand RNA virus and belongs to the family of Betacoronaviruses. Besides hCoV-229E, hCoV-OC43, hCoV-NL63, hCoV-HKU1, SARS-CoV and MERS-CoV, SARS-CoV-2 is the seventh coronavirus known to infect humans^2^. It uses ACE2 receptor for initial attachment to the host cells and, subsequently, requires cleavage of the Spike protein by TMPRSS2 to enter the cells^3,4^. Hence, expression of ACE2 and TMPRSS2 is a pre-requisite for successful infection of cells. Single-cell RNA sequencing studies have provided evidence that these host factors are widely expressed in many organs, highlighting ACE2^+^ TMPRSS^+^ co-expressing cells, such as type 2 alveolar (AT2) epithelial cells, as targets for SARS-CoV-2 infection^5–8^. The most severe manifestation of COVID-19 is observed in the distal lung and manifests as acute respiratory distress syndrome (ARDS)^9^. Diffuse alveolar damage, the underlying histopathologic cause of ARDS, and coagulopathy are the major pathologic findings in lungs of deceased COVID-19 patients. Moreover, activation of the alveolar microvasculature and microthrombosis have been commonly reported in fatal cases of COVID-19^10–12^. In the beginning of the pandemic outbreak about 14% of COVID-19 cases were classified as severe COVID-19^13,14^. Vaccination was an efficient measure to combat SARS-CoV-2 infection and has protected individuals from mild and severe COVID-19^15^. However, due to a decrease of antibody titres over time and viral evolution giving rise to new virus variants that might more efficiently evade the immune system^16^, SARS-CoV-2 infection can still have fatal consequences, particularly in the high-risk population (elderly people with comorbidities or immune deficiency)^9^. Currently, three antiviral compounds – Remdesivir (Gilead Sciences), Molnupiravir (Merck & Ridgeback Biotherapeutics), and Paxlovid (Pfizer) – and five immunomodulatory agents – dexamethasone, Tocilizumab (Roche), Barcitinib (Lilly Eli), Anakinra (Sobi) and Vilobelimab (InflaRX) – are approved for clinical use in COVID-19 patients. However, their applicability is limited to specific patient groups for multiple reasons such as administration in a healthcare setting, interactions with other drugs, concerns to accelerate viral evolution and the emergence of novel virus variants^17^. Therefore, the development of novel targeted therapeutic approaches is still of interest and can be accelerated by using human-derived models, which enable to study human-virus interactions^18^.

The respiratory tract comprises the conducting airways and the alveoli, where gas exchange occurs. In the alveoli, type 1 alveolar (AT1) epithelial cells form a very thin but tight barrier with lung microvascular endothelial cells allowing efficient passive diffusion of oxygen and CO_2_ between air and the blood stream^19,20^. AT2 cells are the other principal constituent of the alveolar epithelium and secrete pulmonary surfactant to prevent the lungs from collapsing during exhalation^21^. Moreover, they are part of the innate immune defence of the alveoli and serve as alveolar progenitor cells. AT2 cells are capable of self-renewal and differentiation into AT1 cells to re-populate the epithelium after lung injury^22^. As a consequence of breathing motion, the alveoli are constantly exposed to mechanical forces (shear stress, surface tension and stretch) which affect key biological functions in lung development^23^, surfactant secretion^24–27^ and wound healing^28–30^. This complex interplay between the different cell types, their microenvironment, and physical forces makes it challenging to model the alveoli *in vitro*. However, recent advances in 3D cell culture have expanded the toolbox of human-derived alveolar models including lung organoids, precision-cut lung slices (PCLS), alveolar 3D microtissues and Lung-on-Chip (LOC) systems^31,32^. The use of these models has been expanded during the COVID-19 outbreak as they recapitulate key aspects of SARS-CoV-2 infection^33–47^. Because they are composed of human cells, there are no uncertainties due to interspecies differences. Therefore, these models offer an opportunity for improved translatability to the clinical setting, aiding in the decision-making process for prioritization of lead compounds.

In this study, we aimed to recreate the air-blood barrier *in vitro* for studying SARS-CoV-2 infection in the alveoli. In our approach, we reduced the complexity of the alveoli to the essential alveolar barrier – formed by alveolar epithelial cells and lung microvasculature endothelial cells – and incorporated essential physical cues such as the air-liquid interface (ALI) and cyclic stretch to recapitulate the epithelial-endothelial barrier under physiological breathing motion. This model recapitulates SARS-CoV-2-induced alveolar damage and was employed for efficacy testing using remdesivir (**Figure 1**).

**Figure 1:**
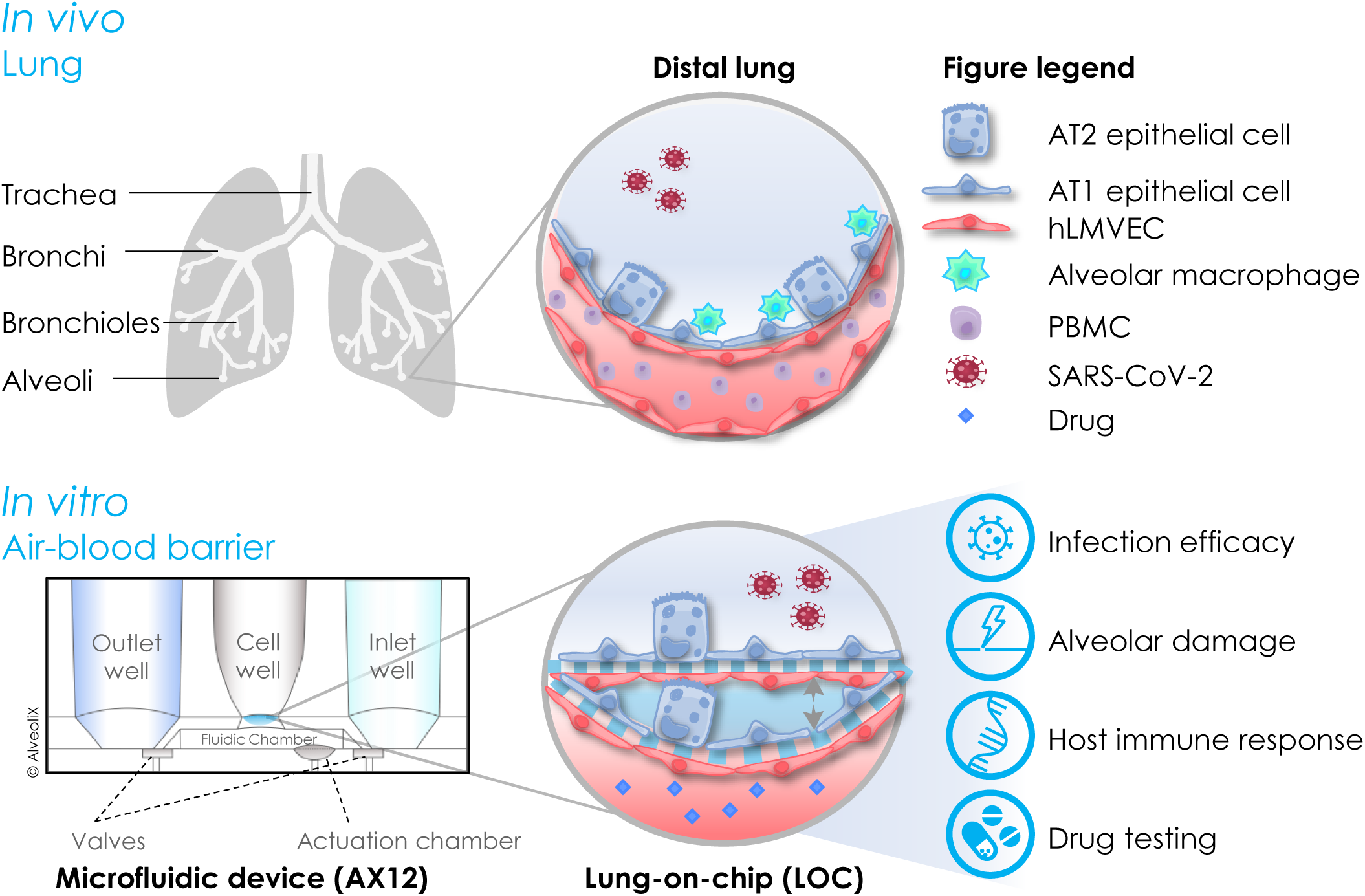
Pre-clinical Lung-on-Chip (LOC) model of SARS-CoV-2 infection. The human respiratory system consists of the upper (trachea, bronchi) and lower (bronchioles, alveoli) airways. In the alveoli, gas exchange occurs through the air-blood barrier, which is comprised of human lung microvascular endothelial cells (hLMVEC), alveolar type 1 (AT1) and alveolar type 2 (AT2) epithelial cells. The alveolar barrier is constantly exposed to the outside environment via inhalation of nanoparticles or pathogens such as SARS-CoV-2. Alveolar macrophages and circulating peripheral blood mononuclear cells (PBMC) exert important functions in mucosal immunity at the alveolar surface. Mechanical forces that stem from breathing motion modulate cell behaviour and function in the distal lung. Therefore, it is challenging to model the alveoli. Advanced *in vitro* models such as Lung-on-Chip (LOC) systems preserve important aspects of the alveolar barrier and can be applied in low-to medium-throughput screens for pre-clinical applications. The AX12 (AlveoliX AG) allows to re-build the alveolar barrier on an ultra-thin, elastic membrane and apply cyclic stretch through the actuation chamber to simulate breathing. We have incorporated hLMVEC, an in-house generated immortalized alveolar epithelial cell line (^AX^iAEC) and cyclic stretch to develop a LOC SARS-CoV-2 infection model and applied the system to study SARS-CoV-2 infection efficacy in the distal lung, evaluated alveolar damage, innate immune responses and assessed anti-viral drug efficacy *in vitro*.

## Material & Methods

### Cells and cell culture

Vero E6 cells were kindly provided by Prof. R. Dijkman (University of Bern) and were cultured in Dulbecco’s Modified Eagles Medium (DMEM; Gibco) containing 4.5g/L glucose, GlutaMAX and 1mM sodium pyruvate, supplemented with 10% heat-inactivated fetal bovine serum (FBS; Sigma-Aldrich), 100 IU/ml penicillin, 100µg/ml streptomycin (1% Pen/Strep; ThermoFisher Scientific), 1% MEM non-essential amino acids (Gibco) and 15mM HEPES (Gibco).

Primary human alveolar type 2 cells (^AX^hAEpC) were provided by AlveoliX AG (Switzerland). The cells were isolated from lung tissue resection obtained with the patient’s informed consent and ethical approval. ^AX^hAEpC were seeded on 24-well cell culture inserts (Falcon Cat. No. 353095 or Corning Cat. No. 3470) and cultured in AX Alveolar Epithelial Medium (^AX^AEM; Alveolix AG). The previously published ^AX^hAEpC cell culture and differentiation protocol^48^ was slightly adapted. Medium was refreshed every 2 – 4 days. For the first few days, ^AX^hAEpC were cultured in submerged conditions with 500μl/well ^AX^AEM on the basal side and 200μl/insert ^AX^AEM on the apical side of the insert. Air-liquid interface (ALI) was established between day 3 – 4 in culture by removing medium from the apical side and leaving 400μl/well ^AX^AEM on the basal side. Some ^AX^hAEpC cultures were maintained in submerged condition for the entire experiment duration to compare infection efficacy to ALI cultures. Immortalized alveolar epithelial cells (^AX^iAEC)^49^ were provided by Alveolix AG and were expanded according to the manufacturer’s guidelines. Cells were seeded in 24-well cell culture inserts (Falcon Cat. No. 353095 or Corning Cat. No. 3470) in AX Alveolar Epithelial Barrier Medium (^AX^AEBM; Alveolix AG) supplemented with 1% Pen/Strep. Cells were maintained submerged in 500µl/well ^AX^AEBM on the basal side and 200µl/insert ^AX^AEBM on the apical side. ALI was established between days 10 – 12, adapting the published protocol for ^AX^iAEC culture in ALI^48,49^. During culture in ALI, cells were maintained in 400µl/well ^AX^AEBM on the basal side.

Human lung microvascular endothelial cells (hLMVEC) were provided by Alveolix AG.

All cell cultures were maintained at 37°C in a humidified incubator with 5% CO_2_.

### Virus

All experiments were performed with SARS-CoV-2, S^D614G^ Wuhan (GenBank: MT108784) at passage 3^50,51^. The virus was kindly provided by Prof. R. Dijkman, University of Bern. The virus construct was originally grown on Vero-TMPRSS2 cells and the virus stock used in this study was produced in Calu-3 cells. Cell culture supernatant from the same batch of uninfected Calu-3 cells was collected in parallel and stored at −80°C as mock control for infection experiments. Viral RNA from the stock was extracted using the QIAamp viral RNA mini kit (Qiagen, Cat. No. 52904) according to the manufacturer’s protocol and genomic stability of the SARS-CoV-2 stock was validated by whole-genome sequencing on a MinION device (Oxford Nanopore Technologies).

Virus stock production and all experiments involving infectious SARS-CoV-2 were conducted in a biosafety level 3 (BSL3) laboratory of the Institute for Infectious Diseases (IFIK), University of Bern, in adherence to Swiss law and international biosafety guidelines (e.g. WHO).

### Alveolar barrier on the Lung-on-Chip (LOC) system

The ^AX^Lung-on-Chip (LOC; Alveolix AG) consists of the LOC, called AX12, two electro-pneumatic units and a connecting interface. The AX12 is connected to the electro-pneumatic units, the ^AX^Exchanger in the cell culture hood, or the ^AX^Breather in the incubator via the ^AX^Dock to control the chip. The AX12 comes in a 96-well plate format and comprises two chips. Each chip contains six cell culture units. One unit consists of an inlet, cell well and outlet, which are connected to each other by basal microfluidic channels and pneumatically controlled valves (**Figure 1**). The cell chamber is divided into basal and apical compartment by a thin (3,5µm thickness), elastic and porous (3µm pore size) membrane.

For the ^AX^iAEC/hLMVEC co-culture model, hLMVEC were seeded on the basal side of the AX12 membrane. Subsequently, the chip was closed, and the basolateral chamber was filled with ^AX^E2-ABM co-culture medium according to the manufactureŕs protocols and as described by Sengupta and colleagues^49^. Finally, ^AX^iAEC were seeded on the apical side of the membrane in ^AX^E2-ABM co-culture medium. Cells were allowed to attach and form a tight barrier in submerged/static conditions until day 8 with regular medium exchanges every second day. Starting time points for ALI and stretch were optimized for the infection experiments in this study. ALI was started on day 8 and stretch was implemented on day 11 in all LOC experiments. The AX12 was placed in the ^AX^Dock in the incubator and connected to the ^AX^Breather. The ^AX^Breather implements 3D cyclic stretch to the cell culture membrane via a microdiaphragm at the bottom of the AX12 plate. ^AX^iAEC/hLMVEC were cultured in ALI/static (non-breathing) or ALI/dynamic (breathing) conditions at 37°C in a humidified incubator with 5% CO_2_.

### SARS-CoV-2 infection on cell culture inserts

^AX^hAEpC were infected between day 7 – 12 from seeding on cell culture inserts. The SARS-CoV-2 inoculum was prepared in 200μl/insert ^AX^AEM to reach an MOI of 0.5. The mock control was prepared in the same manner. ^AX^hAEpC were infected with SARS-CoV-2 inoculum or mock control from the apical side during medium exchange and were incubated at 37°C for 2h to allow virus attachment. Subsequently, the virus inoculum was removed, and cells were twice washed from the apical side with 200μl/insert Dulbecco’s phosphate-buffered saline containing calcium and magnesium (DPBS; ThermoFisher Scientific). Medium exchange was performed on the basal side with 500µl/well or 400µl/well fresh ^AX^AEM for submerged and ALI culture, respectively.

The same protocol with slight adaptations was followed to infect ^AX^iAEC with SARS-CoV-2. Briefly, ^AX^iAEC were infected between day 27 – 33 from seeding on cell culture inserts. ^AX^iAEC cultures were infected either after maturation in ALI or in submerged culture conditions. SARS-CoV-2 stock was diluted in DMEM supplemented with 1% FBS and 1% P/S to reach an MOI of 0.5 in 200µl inoculum. ^AX^iAEC were infected from the apical side and incubated at 37°C. Control inserts were incubated with 200µl/insert mock control on the apical side. After 2h, the inoculum was removed, and cells were washed twice with 200µl DPBS containing calcium and magnesium from the apical side. Medium exchange was performed with fresh ^AX^AEBM and submerged or ALI culture was re-established. The virus inoculum and last DPBS washes at 2h post infection (hpi) were saved and stored at −80°C for virus titration (see section below).

Infected cultures were monitored for cytopathic effect (CPE) at 24hpi, 48hpi and 72hpi by bright field imaging with an Olympus CKX53 microscope (Olympus).

### SARS-CoV-2 infection on the LOC

^AX^iAEC/hLMVEC co-cultures were seeded and differentiated in ALI as described above. On day 11, AX12 were transported to the BSL3 laboratory to start stretch. On day 16, cells were infected from the apical side with SARS-CoV-2 at MOI 0.5. The SARS-CoV-2 inoculum and mock control were prepared in 70µl ^AX^E2-ABM co-culture medium. The AX12 was placed in the ^AX^Dock connected to the ^AX^Exchanger and 70µl/well inoculum was added to the apical side. The medium in inlet and outlet was equilibrated and the AX12 was placed back in the incubator. After incubation at 37°C for 2h, the AX12 was again connected to the ^AX^Exchanger and the inoculum was removed. The apical side was washed twice with 200µl/well DPBS (with calcium and magnesium) and a medium exchange was performed with ^AX^E2-ABM from the basal side according to the manufacturer’s instructions. The last wash from the apical side was kept for virus titration. ALI was re-established in the ^AX^iAEC/hLMVEC co-culture and the AX12 was placed back in a humidified incubator at 37°C and with 5% CO_2_ atmosphere. Infected LOC systems were cultured in ALI/static or ALI/dynamic condition for up to 48hpi and monitored regularly for CPE by bright field microscopy (Olympus CKX53).

### Drug treatment of infected ^AX^iAEC on cell culture inserts

^AX^iAEC were differentiated on cell culture inserts starting ALI culture on day 6 from seeding. On day 14, ^AX^iAEC were infected with SARS-CoV-2 in ^AX^AEBM from the apical side. ^AX^iAEC monocultures were treated with serial dilutions (0.04μM, 0.2μM, 0.5μM, 1μM and 5μM) of remdesivir (MedchemExpress, Cat. No. HY-104077) comparing two protocols. In approach 1, ^AX^iAEC were infected with SARS-CoV-2 at an MOI of 0.5 from the apical side. After 2h, the inoculum was removed and remdesivir treatment was started from the basal and apical side in ^AX^AEBM in submerged condition. Infection efficacy was evaluated at 48hpi. For approach 2, drug treatment was started together with infection at 0hpi. After removing the inoculum at 2hpi, ^AX^iAEC were treated in ALI from 2hpi to 48hpi from the basal side only. In brief, 500μl/well ^AX^AEBM containing remdesivir were added to the basal side and 200μl/insert ^AX^AEBM containing remdesivir and SARS-CoV-2 inoculum at MOI of 0.5 (or mock control) were added to the apical side. After 2h incubation at 37°C, the inoculum was removed from the apical side, cells were washed twice with DPBS (+calcium and +magnesium) and placed back in a humidified incubator at 37°C and a 5% CO_2_ atmosphere. ^AX^iAEC were monitored for CPE and TER, and samples were collected at 48hpi for immunofluorescence staining.

Remdesivir was dissolved in dimethylsulfoxide (DMSO; Sigma-Aldrich). DMSO dilution corresponding to the highest drug concentration used was included as vehicle control (0.5% DMSO).

### Drug treatment of infected LOC

^AX^iAEC/hLMVEC co-cultures on the LOC were infected with SARS-CoV-2 at MOI of 0.5 on day 16 and remdesivir treatment was started simultaneously (at 0hpi), adapting the protocol developed in approach 2 on cell culture inserts. In brief, 1μM Remdesivir or 0.1% DMSO vehicle control were prepared in ^AX^E2-ABM medium and basal medium exchange was performed according to manufacturer’s instructions. The apical inoculum was prepared in 70μl/well ^AX^E2-ABM medium containing remdesivir or vehicle control. Medium was removed from the apical side, and the inoculum was added to the cells for SARS-CoV-2 infection at MOI of 0.5 and simultaneous start of remdesivir treatment. The LOC was incubated at 37°C for 2h, before removing the inoculum from the apical side. ^AX^iAEC were washed twice from the apical side with 200μl/well DPBS (+calcium and +magnesium). ALI was re-established with ^AX^E2-ABM medium containing therapeutic compound (1μM remdesivir or 0.1% DMSO). The AX12 was placed back in the ^AX^Dock and breathing cycles were continued for the next 48h. At 48hpi, TER was measured, and samples were collected for immunofluorescence staining and bulk RNA-sequencing.

### Transepithelial electrical resistance (TER) measurement

Formation of the epithelial barrier was assessed by TER measurement starting from day 2 and thereafter every second day until experiment termination. For ^AX^hAEpC, TER measurements were performed less frequent than for ^AX^iAEC, only every 2 – 4 days. A chopstick electrode (STX2; World Precision Instruments) and a Volt/Ohm meter (EVOM2 or EVOM3; World Precision Instrument) were used to measure TER in cell culture inserts. Briefly, the electrode was sterilized in 70% ethanol and washed in sterile deionized water and DPBS (+calcium and +magnesium) before measurement. 200µl warm DPBS (+calcium and +magnesium) were added to the inserts in ALI culture from the apical side and plates were allowed to equilibrate to room temperature. TER was measured in all cell culture wells and in a background insert without any cells seeded.

TER measurements in the LOC were performed every second day starting from day 2 after seeding until experiment termination as described by Sengupta and colleagues^49^. A 96-well plate electrode (STX100C96, World Precision Instruments) was sterilized in 70% ethanol and washed in sterile deionized water and DPBS (+calcium and +magnesium). The AX12 was placed in the ^AX^Dock under the cell culture hood and the ^AX^Exchanger was started. Wells in ALI were equilibrated to submerged condition by adding warm DPBS containing calcium and magnesium to the apical side and equilibrating the volumes in the inlet and outlet. The LOC was equilibrated to room temperature, then TER measurement was started by selecting the program “TER measurement” on the ^AX^Exchanger. TER was measured in all cell culture wells and background wells by placing the electrode between the central well and outlet well according to the manufacturer’s instructions.

For TER measurement in infected cell culture inserts or LOC, the electrode was shortly dipped into 70% ethanol and sterile deionized water after each measured well to avoid any transfer of infectious virus to the subsequent wells.

To calculate the final TER as Ohm*cm^2^, the resistance of cells (R_cells_) was first calculated by subtracting the resistance of the background well (R_BKGR_) from the total resistance (R_total_) measured in the well:

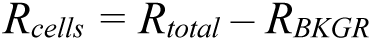

TER was then calculated by multiplying R_cells_ by the surface area (SA) of the insert (0.3cm^2^ or 0.33cm^2^) or AX12 cell culture well (0.071cm^2^):

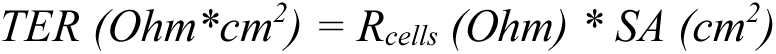

TER values are represented as Ohm*cm^2^ or as fold change relative to the respective experimental control.

### SARS-CoV-2 titration

The supernatant of infected cultures was centrifuged at 500g for 5min to remove dead cells and debris. The cell-free supernatant was transferred to a new tube and stored at −80°C in the BSL3 facility until virus titration. The day before starting virus titration, Vero E6 cells were passaged with Trypsin-EDTA, 0.05% (Gibco), and were seeded in 96-well plates (TPP) at a density of 25’000 – 30’000 cells/well in 125µl/well DMEM supplemented with 10% heat-inactivated FBS, 1% Pen/Strep, 1% MEM non-essential amino acids and 15mM HEPES. The next day, the plates were transported to the BSL3 laboratory and 31µl/well of supernatant containing the virus were added to the top row. Afterwards, 5-fold serial dilutions were performed by transferring 31µl/well from the top row to the next row until the last row was reached. Four wells per condition were used to determine the virus titer. After incubation at 37°C for three days, the plates were fixed in 4% buffered formalin solution (Formafix) and stained with 100µl/well filtered 5% Gram’s crystal violet solution (Sigma, Cat. No. 1092180500) diluted in deionized water. Plates were incubated at room temperature for 30min, then staining solution was removed and plates were washed under running tap water. Wells were evaluated for visible cytopathic effect (CPE), and TCID50/ml was calculated according to the Spearman-Kärber method^52^.

### Immunofluorescence staining and image quantification of cell culture inserts

Infected cell culture inserts were washed with 2x 200µl and 1x 500µl DPBS (+calcium and +magnesium) from the apical and basal side, respectively. The cells were fixed in 500µl/well basal and 300µl/insert apical 4% buffered formalin solution (Formafix). After 10min incubation at room temperature, the inserts were labelled and dunked in 4% buffered formalin solution for transport out of the BSL3 laboratory. After at least 20min additional incubation at room temperature, the inserts were removed from the formalin container, washed in PBS (Sigma; without calcium and magnesium), and stored in PBS at 4°C for immunofluorescence staining.

The membrane was cut out and placed on a glass slide for immunofluorescence staining. Cells were permeabilized in 0.1% Triton X-100 (Sigma-Aldrich, Cat. No. X100) for 10min at room temperature and blocked in 2% bovine serum albumin (BSA; Sigma-Aldrich, Cat. No. A9418) in PBS (2% BSA/PBS) for 1h at room temperature. Primary antibodies were diluted in 2% BSA/PBS blocking solution. After incubation at 4°C overnight, cells were washed in PBS and incubated at room temperature for 90min with 1:200 dilutions of secondary antibodies in 2% BSA/PBS blocking solution. For multiplexing, secondary antibodies were removed and blocked for 1h at room temperature with unconjugated antibody before incubation with pre-conjugated anti-TMPRSS2-AF488 antibody in 2% BSA/PBS blocking solution. Antibodies and dilution used are reported in **Supplementary Table 1**. After incubation at room temperature for 90 minutes, membranes were washed twice with PBS, and nuclear counterstain was performed. Hoechst-33342 (Thermo Scientific, Cat. No. 62249) was diluted 1:1000 in PBS and cells were incubated at room temperature for 10min. Subsequently, membranes were washed twice with PBS and mounted with Fluoroshield^TM^ mounting medium (Sigma-Aldrich, Cat. No. F6182).

For immunofluorescence quantification from cell culture inserts, images were taken with the EVOS M7000 (Life Technologies Corporation) or Leica DMI4000 B (Leica Microsystems) microscope from at least five locations of the cell culture area. Images were analysed in ImageJ 1.52p (Java 1.8.0_172). Background fluorescence was determined from the mean of two to four regions of interest (ROI) per image without positive staining. The background signal was subtracted from the entire image and corrected mean fluorescence intensity (MFI) was measured by imageJ 1.52p. Representative images were acquired with the Leica DMI4000 B (Leica Microsystems) microscope.

### Immunofluorescence staining and image quantification of LOC

Uninfected LOC cultures were fixed in fixative solution (Invitrogen, Cat. No. R37814) for 12 minutes at room temperature as described by Sengupta and colleagues^49^. Afterwards, the LOC was washed with DPBS (without calcium and magnesium) and stored at 4°C until immunofluorescence staining.

Infected LOC cultures were fixed in 4% buffered formalin solution for 20 minutes at room temperature, then washed with DPBS (without calcium and magnesium) and stored at 4°C in the BSL3 laboratory. For transport out of the BSL3 laboratory and immunofluorescence staining, the chip was removed from the plate and disassembled using the ^AX^Disassembly tool (Alveolix AG). The bottom part, containing the cell culture membranes was labelled, and submerged in a container of 4% buffered formalin solution to transport the chips out of the BSL3 facility. After an incubation time of at least 20 minutes at room temperature in 4% buffered formalin solution, the chips were removed from the transport container, washed in PBS and either stored in PBS at 4°C or directly used for immunofluorescence staining.

The disassembled chips were permeabilized in 0.1% Triton X-100 for 10 minutes at room temperature and blocked in 2% BSA/PBS for 1h at room temperature. Staining with primary antibodies was performed at the dilutions indicated in **Supplementary Table 1** in 2% BSA/PBS. The chips were incubated with primary antibody solution at 4°C overnight. The next day, membranes were washed with PBS and incubated for 90 minutes at room temperature with secondary antibodies (**Supplementary Table 1**) in 2% BSA/PBS. For co-staining with conjugated antibodies, the secondary antibodies were blocked for 1h at room temperature with unconjugated primary antibody before incubation with mouse anti-TMPRSS2-AF488 or mouse anti-ZO-1 (**Supplementary Table 1**) diluted in 2% BSA/PBS blocking solution. After incubation at room temperature for 90 minutes, membranes were washed with PBS, and nuclei were stained with Hoechst-33342 used at 1:1000 dilution in PBS. After 10 minutes incubation at room temperature, the chips were washed twice with PBS and were mounted between two cover slips with Fluoroshield^TM^ mounting medium.

To quantify SARS-CoV-2 NP staining and cleaved caspase-3 staining in infected LOC cell culture wells, images were acquired with the EVOS M7000 and stitched to represent the entire well (Life Technologies Corporation). Stitched images were used to determine the corrected MFI of the red (SARS-CoV-2 NP) and far red (cleaved caspase-3) channel by ImageJ 1.52p (Java 1.8.0_172). Briefly, four areas without fluorescence signal were selected from each well to calculate the average background signal per well. The average MFI of the background was subtracted for each well and MFI was measured from the corrected image.

Representative images were acquired by LSM 710 confocal microscope (Carl Zeiss Microscopy).

### RNA isolation and RT-qPCR

For RNA isolation, infected ^AX^iAEC monocultures on cell culture inserts were washed with calcium- and magnesium-free DBPS (Sigma) from the apical and basal side before lysis in 350μl/insert TRI reagent (Zymo Research, Cat. No. R2050). The apical cell lysate was collected in a screw-cap tube (Sarstedt) and transported outside of the BSL3 laboratory after inactivation of infectious virus. Total RNA was isolated using the Direct-zol RNA Microprep kit (Zymo Research, Cat. No. R2063) according to the manufacturer’s instructions. For gene expression analysis, cDNA was synthesized from 500ng RNA by M-MLV reverse transcriptase (Promega, M1701) and qPCR with SYBR^TM^ Select Master Mix (Applied biosystems, Cat. No. 4472908) was run on the TaqMan ViiA^TM^ 7 Real-Time PCR System (Applied Biosystems). Expression of IFNB1, IFNL1, MX1, IL6, ACE2 and TMPRSS2 was analysed and normalized to HPRT-1 and H3F3A housekeeping genes. In addition, viral RNA was detected with specific primers to SARS-CoV-2 N1. Primer sequences were taken from PrimerBank^53^ or CDC repositories as indicated in **Supplementary Table 2**. If no published sequences were available, primers were designed using the NCBI online tool PrimerBlast^54^. All primers were tested *in silico* (UCSC In-Silico PCR online tool of UCSC Genome Browser^55^) and validated in ^AX^iAEC before use.

### Sample preparation for bulk RNA-sequencing

^AX^iAEC RNA from infected LOC co-cultures was sent for bulk RNA sequencing. Medium exchange with DPBS (without calcium and magnesium) was performed according to the manufacturer’s instructions for cell lysis. Subsequently, ^AX^iAEC were washed with calcium- and magnesium-free DBPS and lysed from the apical side in 200μl/well TRI reagent. Sample tubes were sterilized and transported out of the BSL3 laboratory for RNA isolation using the Direct-zol RNA Microprep kit (Zymo Research, Cat. No. R2063). ^AX^iAEC RNA quality was determined for each sample by Qubit^TM^ 4 Fluorometer (Thermofisher Scientific) using the Qubit^TM^ RNA Integrity and Quality (IQ) Assay kit (Thermofisher Scientific, Cat. No. Q33221) and by 2100 Bioanalyzer (Agilent) using the RNA 6000 Nano kit (Agilent, Cat. No. 5067-1511) or RNA 6000 Pico kit (Agilent, Cat. No. 5067-1513) depending on the RNA amount available. Samples with a RIN > 7.0 passed minimal quality requirements and were prepared for bulk RNA sequencing. The RNA concentration was determined by Qubit^TM^ RNA Braod Range (BR) Assay kit (Thermofisher Scientific, Cat. No. Q10210) on a Qubit^TM^ 4 Fluorometer. Samples were diluted to an RNA concentration of 20ng/μl and 200ng – 400ng RNA were shipped for bulk RNA-sequencing.

### Bulk RNA-sequencing

Library preparation and sequencing was performed in Alithea Genomics SA (Lausanne, Switzerland) using multiplexed 3’-end bulk RNA barcoding and sequencing (MERCURIUS^TM^ BRB-seq service). Read trimming, alignment, and quantification steps were performed by Alithea Genomics using STARsolo^56^ and the GRCh38.104 (human) reference genome. UMI-deduplicated count matrices were used for the downstream analyses utilizing *DESeq2* (version 1.46.0). Technical replicates were collapsed using the *collapseReplicates* function. Differentially expressed genes (DEGs) were selected with an adjusted p-value cutoff of 0.01. Pathway analysis was performed using *decoupleR*^57^ and the PROGENy pathways^58^.

### Statistics

Outliers were identified in pooled replicates by GraphPad Prism’s online outlier calculator (https://www.graphpad.com/quickcalcs/grubbs1/) assuming α = 0.05. Outliers were removed from analysis. Normal distribution of replicates was checked in GraphPad Prism version 10.4.1 (GraphPad Prism Software) and means of each independent experiment (N) were calculated. The data was plotted as XY plots showing mean ± SD or as superplots in GraphPad Prism version 10.4.1. Superplots display mean ± SD of N independent experiments and distribution of n technical replicates in an overlaid graph^59^. Statistics was performed on the mean of minimum two independent experiments to evaluate differences between groups. The specific statistical tests are indicated in the figure legends. All statistical tests were performed with a significance level of α = 0.05 and p values lower than 0.05 were considered as statistically significant (*, p ≤ 0.05; **, p ≤ 0.01; ***, p ≤ 0.001; ****, p ≤ 0.0001).

## Results

### SARS-CoV-2 infection of alveolar epithelial cells in air-liquid interface (ALI) culture

Since multiple studies have reported SARS-CoV-2 infection in the alveolar epithelium^8,60^, we first evaluated primary human alveolar epithelial cells (^AX^hAEpC) to build the SARS-CoV-2 infection model. ^AX^hAEpC have previously been characterized for *in vitro* toxicity studies^48^. To test their susceptibility to SARS-CoV-2 infection, ^AX^hAEpC were isolated from lung resection tissue of four donors (**Supplementary Table 3**) and were differentiated in ALI on cell culture inserts. Whenever possible, we evaluated expression of SP-C, a marker of AT2 cells, and host factors for SARS-CoV-2 infection. While SP-C^+^ cells were present in all cultures assessed, the expression of ACE2 and TMPRSS2 was variable among different donors (**Supplementary Figure 1A**). The barrier formation capacity was evaluated as an additional measure of ^AX^hAEpC quality. Two out of four donors (donor 2 and donor 3) did not form a stable alveolar barrier in ALI culture and never reached TER values above 200 Ohm*cm^2^. ^AX^hAEpC from donor 1 and donor 4 formed a tight barrier in ALI culture (**Supplementary Figure 1B**). Next, we quantified SARS-CoV-2 titers in the apical washes of ^AX^hAEpC ALI cultures and found that SARS-CoV-2 infection was abortive in ^AX^hAEpCs of donor 2 and donor 3 (**Supplementary Figure 1C**). In contrast, SARS-CoV-2 progeny was released from ^AX^hAEpCs of donor 1 and donor 4, however, with differential efficacy (**Supplementary Figure 1C**). SARS-CoV-2 titers in ^AX^hAEpC cultures from donor 1 increased more than two logs from 1.33 x 10^3^ TCID50/ml at 2hpi and 2.54 x 10^4^ TCID50/ml at 48hpi up to 1.735 x 10^5^ TCID50/ml at 72hpi (p ≤ 0.001). Interestingly, donor 1, was diagnosed with COPD (**Supplementary Table 3**), which increases the risk of severe COVID-19^61^. However, our patient cohort is too small to draw any conclusion on SARS-CoV-2 susceptibility and COPD. A weaker but still significant 3-fold increase of SARS-CoV-2 titers was reached in ^AX^hAEpC cultures from donor 4, increasing from 1.28 x 10^2^ TCID50/ml at 2hpi and 2.93 x 10^2^ TCID50/ml at 48hpi to 3.88 x 10^2^ TCID50/ml at 72hpi (p ≤ 0.05).

Interestingly, IF staining for SARS-CoV-2 nucleocapsid protein (NP) in infected ^AX^hAEpC ALI cultures at 48hpi and 72hpi revealed, that infected cells were present in all four donors at both time points (**Supplementary Figure 2A & C**) despite the absence of viral titers in the supernatant of ^AX^hAEpCs of donor 2 and 3. Notably, SARS-CoV-2 infection in submerged conditions was less efficient than in ALI culture. Only few or no SARS-CoV-2 NP^+ AX^hAEpC were observed in submerged cell culture (**Supplementary Figure 2B & D**). Interestingly, ^AX^hAEpCs of donor 4 were susceptible to SARS-CoV-2 infection and produced viral progeny in ALI culture, while SARS-CoV-2 NP was no longer detectable in submerged cultures at 72hpi (**Supplementary Figure 2D**). In line, significantly lower SARS-CoV-2 titers were released from submerged cultures compared to ALI cultures at 72hpi (**Supplementary Figure 2E,** p ≤ 0.05).

These data indicate that alveolar epithelial cells are more susceptible to SARS-CoV-2 infection in ALI. To have a stable cell source and circumvent donor-to-donor variability at the drug testing stage, we leveraged an ^AX^hAEpC-derived alveolar epithelial cell line, ^AX^iAEC, for SARS-CoV-2 infections. This immortalized cell line consists of a mixture of AT1 and AT2 cells^49^. ^AX^iAEC express ACE2 and TMPRSS2^49^, which are exploited by SARS-CoV-2 for initial cell attachment and subsequent cleavage of the Spike protein to enter lung cells^3^. We first assessed ^AX^iAEC susceptibility to SARS-CoV-2 infection on cell culture inserts to evaluate the effect of ALI culture on virus production and alveolar barrier damage (N = 2 experiments). Interestingly, ALI culture induced a slight but not significant upregulation of ACE2 mRNA (**Supplementary Figure 3A**) paralleled by a more efficient SARS-CoV-2 infection in ALI than submerged culture. This was consistently supported by IF staining for SARS-CoV-2 NP (**Supplementary Figure 3B & C**), virus titration (**Supplementary Figure 3D – H,** p ≤ 0.01 at 48hpi) and TER measurements (**Supplementary Figure 3I – L,** p ≤ 0.001 at 48hpi, p ≤ 0.01 at 72hpi). In ALI culture, SARS-CoV-2 infection peaked at 48hpi, showing infection sites corresponding to large areas of the ^AX^iAEC cultures, releasing the highest observed SARS-CoV-2 titers to the apical side (64’183 TCID50/ml), and exerting the strongest impact on barrier integrity (approx. 80% reduction of TER to 532 Ohm*cm^2^). In line with these findings, SARS-CoV-2 replication and interferon (IFN) response peaked at 48hpi in ALI cultures (**Supplementary Figure 4A - C**). The highest upregulation was measured for IFNL1 increasing by 5’421-fold in mRNA levels in infected ALI cultures at 48hpi (**Supplementary Figure 4C,** p ≤ 0.01). In contrast to IFNs, mRNA levels of the interferon-stimulated gene (ISG), MX1, and inflammatory cytokine IL6 were not downregulated after the peak at 48hpi but remained highly expressed at 72hpi (**Supplementary Figure 4D & E**), again supporting a stronger upregulation observed in ALI than in submerged cultures.

These results highlight the importance of alveolar epithelial cell differentiation in ALI to achieve efficient SARS-CoV-2 infection. In ALI, SARS-CoV-2 production and IFN response peaks at 48hpi and disrupts barrier function of infected ^AX^iAEC. Interestingly, the infected ALI cultures show barrier function recovery at 72hpi. Multiple studies reported endothelial cells and stretch as essential modulators of alveolar inflammatory processes and barrier permeability^48,62–66^. We, therefore, aimed to increase the complexity of the model and establish a co-culture of ^AX^iAEC and human lung microvascular endothelial cells (hLMVEC) in a more physiological environment including stretch by leveraging a LOC system.

### The LOC model recapitulates the alveoli under physiological stretch

We seeded ^AX^iAEC on the apical side and human lung microvascular endothelial cells (hLMVEC) on the basal side of the membrane to recreate the epithelial/endothelial barrier of the alveoli in the LOC. ALI was implemented on day 8, and stretch was started on day 11 after seeding. Cyclic 3D stretch was exerted on the elastic cell culture membrane by pneumatic forces controlled by the ^AX^Breather. By day 16, the TER values in the stretched LOC stabilized and the co-cultures were used for infections and downstream experiments (**Figure 2A**). In static cultures, the TER of ^AX^iAEC/hLMVEC co-cultures reached 1700 Ohm*cm^2^ by day 16 (**Figure 2B**). ALI/dynamic cultures stabilised after starting stretch (**Figure 2C**) and ^AX^iAEC/hLMVEC co-cultures formed a tight barrier at 630 Ohm*cm^2^ (**Figure 2B**). We further assessed the expression of AT1 and AT2 cell markers by immunofluorescence staining in ALI/static and ALI/dynamic cultures. We did not observe major differences in the expression of AT1 and AT2 markers and confirmed the presence of AT1 cells (HTI-56) and AT2 cells (SP-C) in both co-culture systems by immunofluorescence staining (**Figure 2D**). Moreover, we further confirmed the tight epithelial/endothelial barrier in static and dynamic conditions, as indicated by the expression of ZO-1 in the epithelial compartment and VE-CAD in the endothelial compartment (**Figure 2D**). Regarding infection experiments, we confirmed the expression of ACE2 and TMPRSS2 in our LOC model (**Figure 2D**).

**Figure 2:**
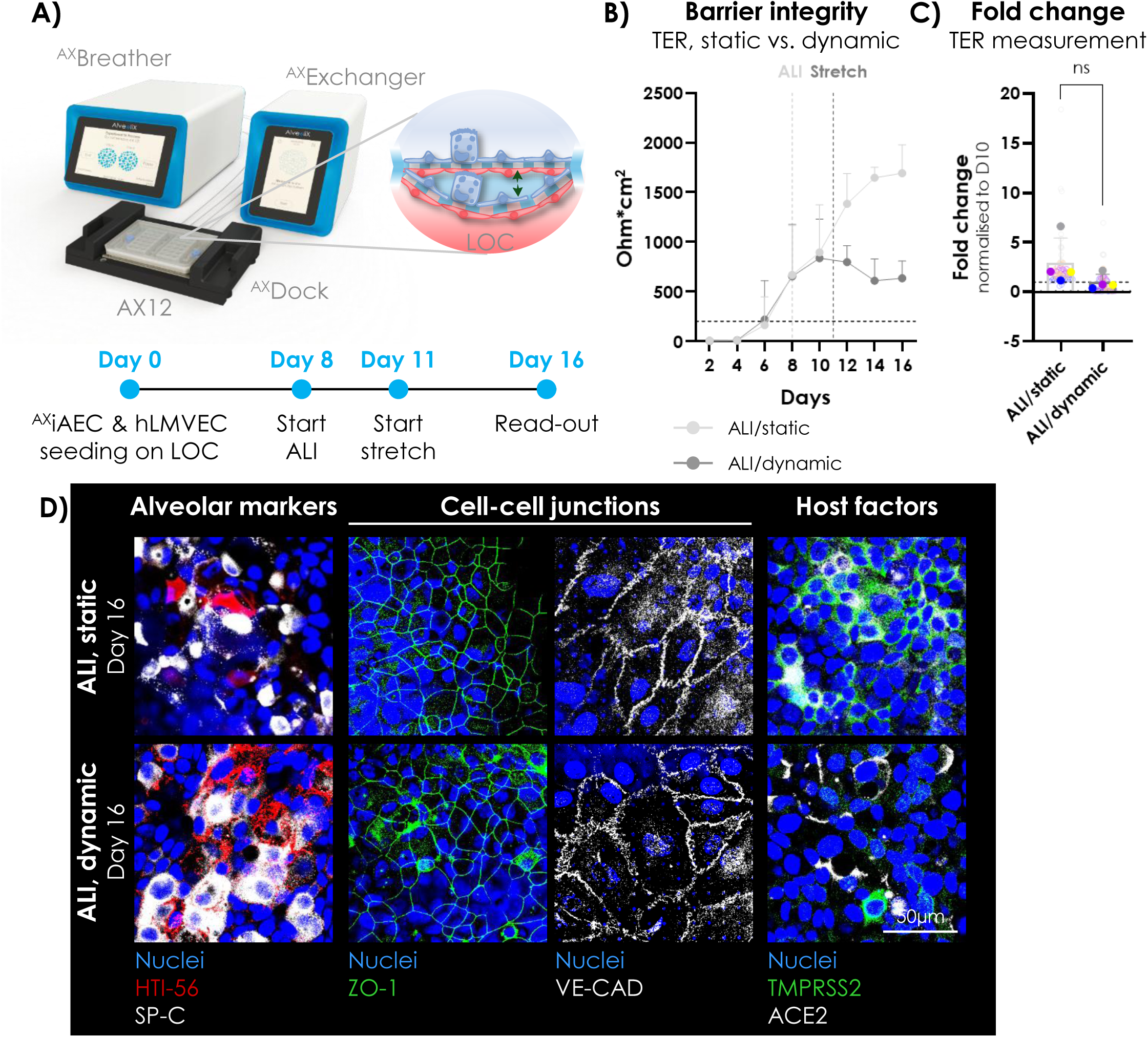
The LOC recapitulates the alveolar barrier under physiological stretch. **(A)** Schematic of the LOC setup. The alveolar barrier is recreated in the AX12 by ^AX^iAEC on the apical side and hLMVEC on the basal side of an ultra-thin and soft, porous membrane. The ^AX^Dock connects the AX12 to the ^AX^Exchanger for manipulation of the cells (e.g. medium exchange, TER measurement) or the ^AX^Breather to implement breathing motion. The cell maturation protocol was adapted for optimal starting time point of ALI and stretch. After 16 days in culture, the maturation status of the LOC was assessed. **(B)** ALI was started on day 8 (dotted line, light grey) and stretch was implemented on day 11 (dotted line, dark grey) for the LOC in dynamic condition. The barrier was considered as tight when reaching TER values above a threshold of ≥200 Ohm*cm^2^ (dotted line, black). N = 4 independent experiments; n = 10 – 24 replicates per experiment. **(C)** Relative TER values suggest a stagnation of TER after implementing stretch, while TER continues to increase between day 10 to 16 in static condition. n = 66 – 70 well replicates from 6 chips per condition run in N = 4 independent experiments. Statistics: Wilcoxon matched-pairs signed rank test. ns, p > 0.05. **(D)** Expression and location of tight junction markers, AT1 and AT2 markers and SARS-CoV-2 host factors were evaluated in ^AX^iAEC/hLMVEC co-cultures after 16 days in culture under ALI/static or ALI/dynamic condition. HTI-56^+^ (red) AT1 cells and SP-C^+^ (grey) AT2 cells were observed under static and dynamic conditions. Co-cultures formed a barrier as indicated by localization of the tight junction protein ZO-1 (green) and the adherens junction protein VE-CAD (grey) to cell-cell junctions of ^AX^iAEC and hLMVEC, respectively. ACE2 (grey) and TMPRSS2 (green) expression in ^AX^iAEC was confirmed. Representative images are shown. Scale bar = 50µm.

We next assessed how stretch influenced SARS-CoV-2 infection and virus-mediated cytotoxicity in our model system.

### The LOC supports SARS-CoV-2 infection and enables studies on virus-mediated alveolar damage under physiological stretch

^AX^iAEC/hLMVEC co-cultures were matured in ALI starting from day 8 and afterwards either kept in static condition or stretched from day 11 onwards. Cells were infected on day 16 with SARS-CoV-2 at MOI 0.5 under static or dynamic conditions. Infection efficacy, virus production, cytopathic effect (CPE), and cellular response to the virus were evaluated at 48hpi (**Figure 3A**), which is the time point when SARS-CoV-2 infection and alveolar damage peaked in ^AX^iAEC in previous experiments on cell culture inserts (**Supplementary Figures 3 & 4**). CPE manifested as irregular dark patches of cells that were not present in the mock control (**Figure 3B & Supplementary Figure 5A**). Static cultures showed only few areas of syncytial cells, whereas, in dynamic cultures, larger areas of syncytial cells were dispersed among the cell culture (**Figure 3B**), suggesting that cells under stretch were more sensitive to SARS-CoV-2 infection. In addition, bulk RNA-sequencing data of ^AX^iAEC infected in ALI/static or ALI/dynamic suggest that stretch modulates ^AX^iAEC response to SARS-CoV-2 infection. Principal component analysis (PCA) showed that samples grouped according to SARS-CoV-2 infection state (PC1) and the presence of stretch (PC2) as the two parameters explaining most of the variance (**Figure 3C**). SARS-CoV-2 infection of alveolar epithelial cells has been reported to be unproductive in microphysiological systems^39^. In line, our LOC model produced low viral titers. Nevertheless, we consistently detected SARS-CoV-2 progeny released to the apical side of ^AX^iAEC/hLMVEC co-cultures under ALI/static and ALI/dynamic conditions. In static conditions, apical viral titers reached a maximum of 1’725 TCID50/ml at 48hpi, while dynamic cultures released significantly higher viral titers reaching 4’258 TCID50/ml (**Figure 3D**, p ≤ 0.05, N = 2 experiments). Next, we confirmed the presence of infected cells in our LOC model under static and dynamic conditions by IF staining for SARS-CoV-2 NP. Moreover, we stained for cleaved caspase-3 (cl. casp-3) to evaluate SARS-CoV-2-induced apoptosis. We observed few dispersed cl. casp-3^+^ cells in mock control wells (**Supplementary Figure 5B**) but mostly they accumulated in infected areas and co-localized with SARS-CoV-2 NP in infected wells (**Figure 3E**). Quantification of the IF signal indicated an increase of SARS-CoV-2 NP signal compared to mock, though not reaching significance, probably due to the limited number of replicates (**Figure 3F**, N = 2 experiments). We did not observe a difference in cl. casp-3 staining between infected and mock control wells in ALI/static (**Figure 3G**). In contrast, there was a trend for increased cl. casp-3 staining in infected ALI/dynamic cultures compared to mock controls at 48hpi (**Figure 3G**). We observed the most pronounced effect of infection on barrier integrity, particularly in dynamic conditions. In static cultures, SARS-CoV-2 infection decreased absolute TER values at 48hpi (**Figure 3H**) but did not induce a significant reduction of TER compared to mock (**Figure 3I**, N = 2 experiments). Notably, TER remained above 700 Ohm*cm^2^ at all time points measured in infected wells in ALI/static. This indicates that SARS-CoV-2 marginally affects barrier function in ALI/static. In contrast, SARS-CoV-2 infection reduced TER values below 100 Ohm*cm^2^ in dynamic conditions at 48hpi (**Figure 3H**) and induced a nearly 90% loss of barrier function compared to mock control under stretch (**Figure 3I**, p ≤ 0.05, N = 2 experiments).

**Figure 3:**
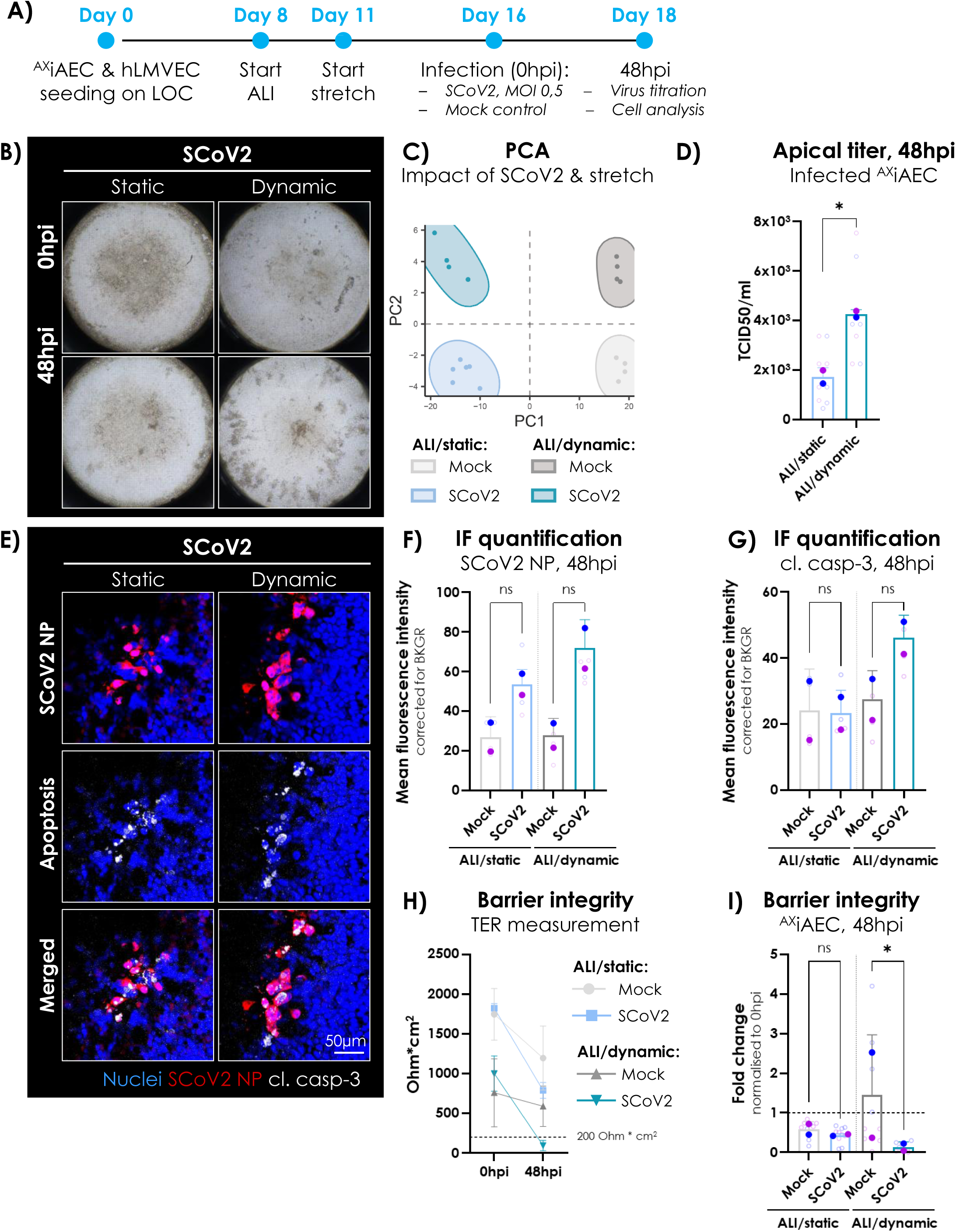
SARS-CoV-2 infects alveolar co-cultures under physiological stretch and allows to study virus-mediated lung damage. **(A)** SARS-CoV-2 infection protocol on the LOC. ^AX^iAEC/hLMVEC co-cultures were matured in ALI/static or ALI/dynamic for 16 days before infection with SARS-CoV-2 at MOI 0.5. Infection efficacy, virus production, barrier tightness and alveolar damage were evaluated at 48hpi. **(B)** Representative images of infected wells at 0hpi and 48hpi in ALI/static and ALI/dynamic. **(C)** RNA was harvested from ^AX^iAEC cells of mock and SARS-CoV-2 infected wells in ALI/static or ALI/dynamic at 48hpi. PCA was performed on transcriptomic data of n = 4 – 6 well replicates from 2 chips per condition run in N = 2 independent experiments. **(D)** TCID50/ml was calculated from apical washes of infected ^AX^iAEC in ALI/dynamic and ALI/static at 48hpi. n = 7 – 10 well replicates from 3 – 4 chips per condition run in N = 2 independent experiments. Statistics: Paired t-test. *, p ≤ 0.05. **(E)** Representative images of ^AX^iAECs infected with SARS-CoV-2 at an MOI of 0.5 in ALI/static or ALI/dynamic culture and stained for SARS-CoV-2 (SCoV2) NP (red) or cl. casp-3 (grey). Nuclei were counterstained with Hoechst-33342 (blue). Cells were fixed and stained at 48hpi. Scale bar = 50µm. **(F) & (G)** Quantification of corrected mean fluorescence intensity (MFI) for SARS-CoV-2 (SCoV2) NP and cl. casp-3 in infected and mock control wells in ALI/static and ALI/dynamic at 48hpi. n = 4 – 5 well replicates from 2 chips per condition run in N = 2 independent experiments. Statistics: Ratio paired t-test. ns, p > 0.05. BKGR = background. **(H)** TER measurement in mock control and SARS-CoV-2 infected wells reveals a drop of TER values in infected wells at 48hpi. A TER of 200 Ohm*cm^2^ was defined as threshold for a tight barrier (dotted line, black). N = 2 independent experiments; n = 4 – 12 replicates per experiment. **(I)** Relative TER values at 48hpi are represented as fold change to 0hpi and the impact of SARS-CoV-2 infection on barrier integrity was evaluated in ALI/static and ALI/dynamic condition. n = 9 – 12 well replicates from 4 chips per condition run in N = 2 independent experiments. Statistics: Ratio paired t-test. *, p ≤ 0.05. SCoV2 = SARS-CoV-2.

In conclusion, SARS-CoV-2 infection damages ^AX^iAEC/hLMVEC co-cultures more drastically under physiological stretch and induces complete barrier breakdown at 48hpi. Given the stronger impact of SARS-CoV-2 infection on the alveolar barrier under stretch, we chose to develop a drug treatment protocol in ALI/dynamic conditions.

### Establishment of a drug treatment protocol in infected ^AX^iAEC

We selected remdesivir for protocol optimization, which is FDA-approved for the treatment of COVID-19 patients^17^ and possesses anti-SARS-CoV-2 activity *in vitro*^43–45^. At first, ^AX^iAEC monocultures were differentiated in ALI from day 6 to day 14 on cell culture inserts. On day 14, ^AX^iAECs were infected with SARS-CoV-2 at an MOI 0.5 to compare two protocols for drug administration route and dose finding. In protocol 1, the cells were incubated with SARS-CoV-2 inoculum on the apical side for 2h. Subsequently, the virus was removed, and drug treatment was started from the basal and apical side at 2hpi. ^AX^iAEC were cultured in submerged condition during the drug treatment to allow a harsher drug treatment schedule with cells being exposed to the drug from the basal and apical side (**Supplementary Figure 6A**). In protocol 2, remdesivir treatment from the apical and basal side was started simultaneously with SARS-CoV-2 infection. After 2h incubation, the inoculum was removed, ALI was re-established. and remdesivir treatment was continued up to 48hpi from the basal side only (**Supplementary Figure 6B**). As expected, we observed patches of syncytial cells in cultures treated with vehicle control according to protocol 2. However, CPE was not observed in cultures that were infected according to protocol 1 (**Supplementary Figure 6C**). Moreover, TER values decreased in cultures that were infected and treated in ALI (protocol 2) but not in cultures that were infected according to protocol 1 suggesting that SARS-CoV-2 infection did not affect ^AX^iAEC when infected and treated in submerged condition from 0hpi to 48hpi (**Supplementary Figure 6D**). Therefore, we could only evaluate remdesivir efficacy in ^AX^iAEC cultures treated according to protocol 2. SARS-CoV-2 infection resulted in a significant decrease of barrier function in infected ^AX^iAEC left untreated or treated with vehicle control (**Supplementary Figure 6E,** p ≤ 0.01 for untreated mock vs. SARS-CoV-2; p ≤ 0.05 for vehicle mock vs. SARS-CoV-2, N = 2 experiments). Remdesivir was effective in the micromolar range resulting in a significant increase of barrier function at 0.2µM (p ≤ 0.01), 0.5µM (p ≤ 0.001) and 1µM (p ≤ 0.01) compared to vehicle control inducing complete recovery of barrier function (**Supplementary Figure 6E**). Remdesivir is an antiviral drug that inhibits the viral RNA-dependent RNA polymerase (RdRp), thereby limiting viral replication^67^. Accordingly, staining for SARS-CoV-2 NP was reduced in a dose-dependent manner in infected cultures and was nearly absent upon treatment with 1µM remdesivir (**Supplementary Figure 7A**). SARS-CoV-2 infection induced a significant increase in SARS-CoV-2 NP signal in infected, untreated cultures and vehicle control (p ≤ 0.0001). Upon remdesivir treatment the signal for SARS-CoV-2 NP staining was strongly reduced at 0.2µM (p ≤ 0.001), 0.5µM (p ≤ 0.001), 1µM (p ≤ 0.0001) and 5µM (p ≤ 0.0001) concentrations (**Supplementary Figure 7B,** N = 2 experiments). Quantification of the signal for cl. casp-3 in mock controls suggested no cytotoxicity of remdesivir in the treatment range of 0.04µM to 5µM (**Supplementary Figure 7C, left**). In infected ^AX^iAEC, we noticed a trend for lower cl. casp-3 signal upon remdesivir treatment compared to vehicle control, but the differences did not reach significance (**Supplementary Figure 7C, right,** N = 2 experiments).

In summary, the drug treatment protocol in ALI (protocol 2) recapitulated the protective effect of the FDA-approved drug remdesivir in ^AX^iAEC monocultures on cell culture inserts. Therefore, we translated this protocol to the LOC infection model and tested efficacy of 1µM remdesivir treatment against SARS-CoV-2 infection in the ^AX^iAEC/hLMVEC co-culture under stretch.

### Treatment of the LOC infection model with the antiviral drug remdesivir

^AX^iAEC and hLMVEC were seeded in co-culture on the LOC and differentiated in ALI starting from day 8. Stretch was implemented on day 11. On day 16, ^AX^iAEC were infected with SARS-CoV-2 at MOI 0.5 from the apical side. At the same time, 1µM remdesivir was administered from the basal and apical side. At 2hpi, the inoculum was removed from the apical side, and the treatment was continued from the basal side in ALI/dynamic up to 48hpi (**Figure 4A**). Examination for CPE at 48hpi revealed more widespread areas of syncytial cells in infected cultures treated with vehicle control compared to infected cultures treated with remdesivir (**Figure 4B, top panel**). IF staining for SARS-CoV-2 NP and cl. casp-3 suggested that remdesivir treatment was highly effective in inhibiting SARS-CoV-2 propagation and limited apoptosis (**Figure 4B, middle and bottom panel**). IF quantification confirmed that SARS-CoV-2 NP signal was significantly increased in infected cultures compared to mock control (p ≤ 0.01) and was significantly reduced by remdesivir treatment (**Figure 4C**, p ≤ 0.05, N = 4 experiments). Moreover, SARS-CoV-2 infection induced a significant increase of cl. casp-3 staining compared to mock control (p ≤ 0.01, N = 4 experiments). Remdesivir treatment slightly decreased apoptosis in infected cultures but was not sufficient to induce a significant decrease in the cl. casp-3 signal during 48h treatment (**Figure 4C**). Finally, barrier function of ^AX^iAEC/hLMVEC co-cultures was significantly impaired upon SARS-CoV-2 infection (p ≤ 0.01) but was significantly increased and restored at 48hpi upon treatment with 1µM remdesivir (p ≤ 0.05, N = 4 experiments; **Figure 4D**).

**Figure 4:**
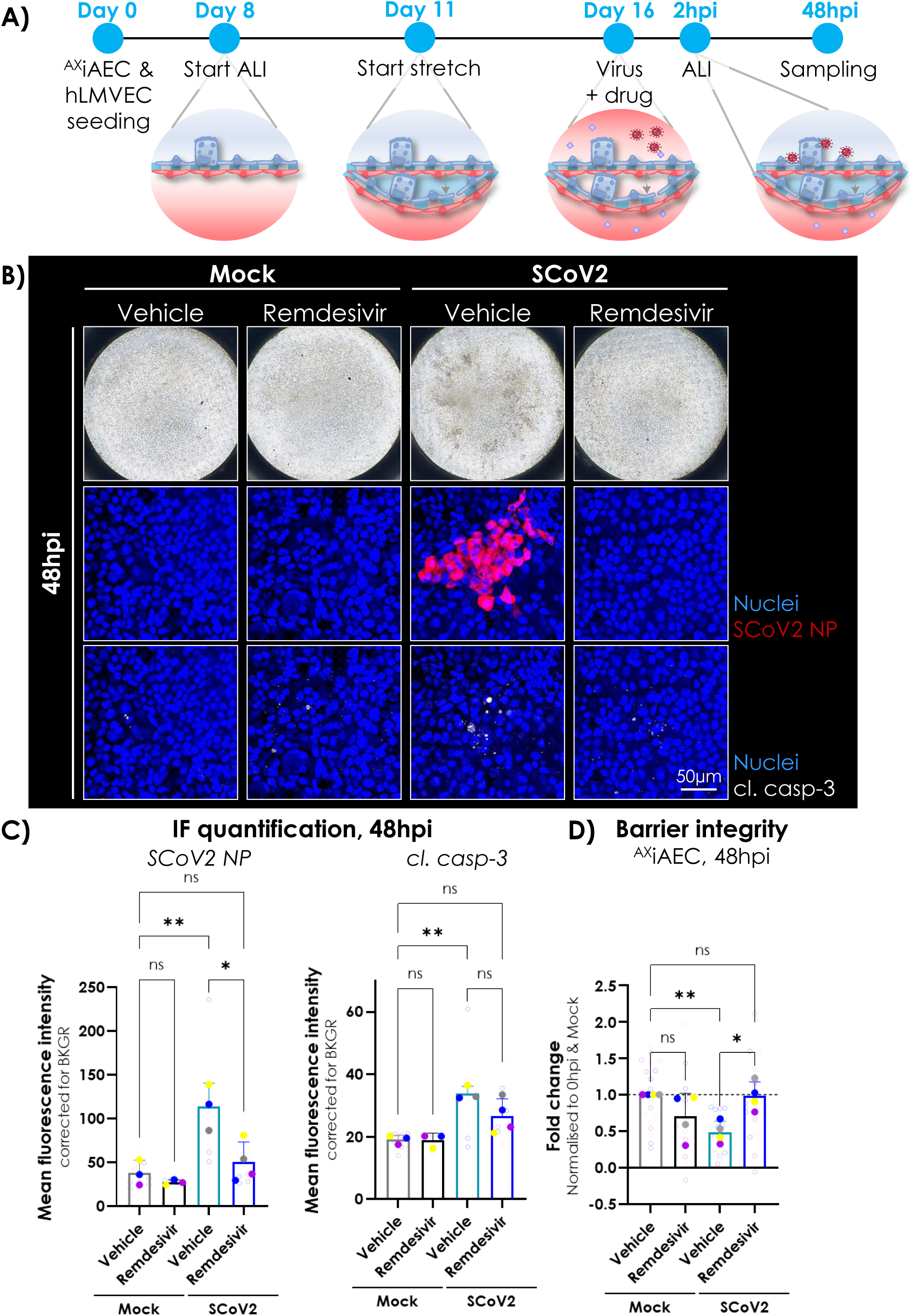
Remdesivir treatment in the LOC infection model. **(A)** SARS-CoV-2 infection and treatment protocol on the LOC. ^AX^iAEC/hLMVEC co-cultures were matured in ALI/dynamic for 16 days before infection with SARS-CoV-2 at MOI 0.5. Treatment with 1µM remdesivir was started at the same time from the basal and apical side. After 2h incubation, the inoculum was removed from the apical side and ALI was re-established. Infected co-cultures were treated with 1 µM remdesivir from the basal side for 48h and, subsequently, were sampled to evaluate SARS-CoV-2 infection efficacy, barrier tightness and alveolar damage. **(B)** *Top panel*: ^AX^iAEC/hLMVEC co-cultures were infected with SARS-CoV-2 or mock control and received treatment with 1µM remdesivir or vehicle control (0.1% DMSO) for 48h in ALI/dynamic. The LOC was inspected for cytopathic effect (CPE) at 48hpi. Images of representative wells are shown. **(B)** *Middle and bottom panel*: Cells in the LOC model were fixed at 48hpi and infection efficacy was evaluated by IF staining for SARS-CoV-2 (SCoV2) NP (red; middle panel). Cell death was evaluated by staining for apoptosis marker cl. casp-3 (grey; bottom panel) and nuclei were stained with Hoechst-33342 (blue). Representative images are shown. Scale bar = 50µm. **(C)** Quantification of corrected mean fluorescence intensity (MFI) for SCoV2 NP (left) and cl. casp-3 (right) in infected and mock control wells with or without 1µM remdesivir treatment in ALI/dynamic at 48hpi. n = 5 – 9 well replicates from 3 – 4 chips per condition run in N = 4 independent experiments. Statistics: Mixed-effects analysis without Geisser-Greenhouse correction and Sidak’s multiple comparison (with a single pooled variance) post hoc test. *, p ≤ 0.05; **, p ≤ 0.01. BKGR = background. **(D)** Relative TER values at 48hpi are represented as fold change to 0hpi and Mock/vehicle control. The impact of SARS-CoV-2 infection on barrier integrity was evaluated at 48hpi after treatment with vehicle control or remdesivir. n = 18 – 23 well replicates from 8 chips per condition run in N = 4 independent experiments. Statistics: Repeated-measures one-way ANOVA without Geisser-Greenhouse correction and Sidak’s multiple comparison (with a single pooled variance) post hoc test. *, p ≤ 0.05; **, p ≤ 0.01. SCoV2 = SARS-CoV-2.

Together these data suggest that remdesivir limits SARS-CoV-2-mediated alveolar damage and is effective to inhibit SARS-CoV-2 replication. Therefore, we next asked, if these changes are also reflected on transcriptional level.

### Remdesivir treatment partially reverts the SARS-CoV-2-mediated transcriptional changes towards a transcriptional profile of uninfected ^AX^iAEC on the LOC

We isolated RNA from all drug treatment experiments on the LOC (N = 4) and performed bulk RNA-sequencing of infected ^AX^iAEC treated with vehicle control or remdesivir and compared the transcriptional profile to uninfected ^AX^iAEC treated with vehicle control. PCA of these three groups revealed that vehicle-treated infected and uninfected ^AX^iAEC grouped separately in the PC1 dimension while infected remdesivir-treated ^AX^iAEC formed a cluster in the middle. They partially overlapped with the mock control suggesting that remdesivir treatment partially reverted SARS-CoV-2-mediated transcriptional dysregulation in infected ^AX^iAEC (**Figure 5A**).

**Figure 5:**
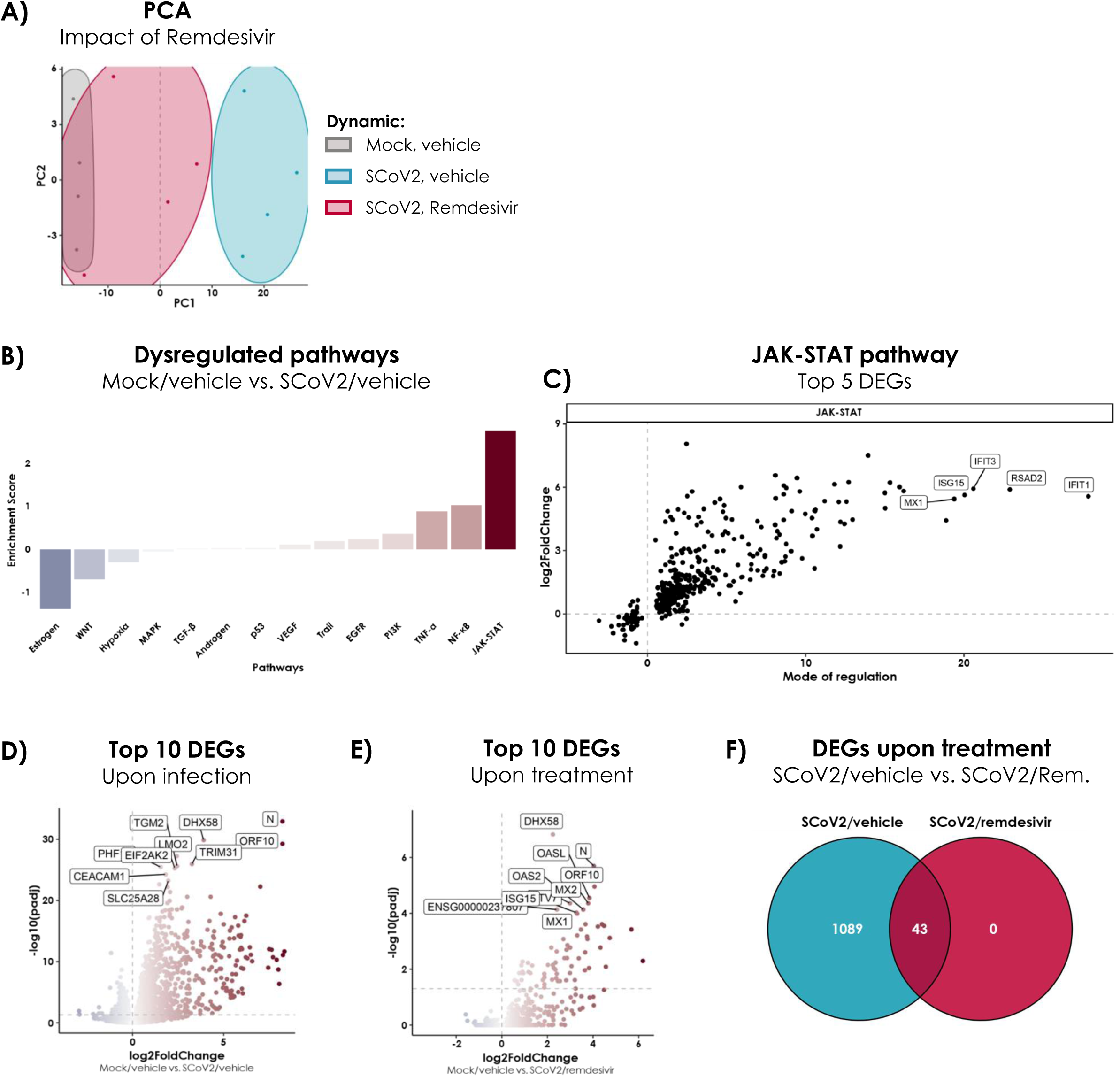
Remdesivir treatment partially restores the transcriptional profile of infected ^AX^iAEC towards a phenotype of uninfected cells. **(A)** PCA reveals a clear separation of infected (SARS-CoV-2/vehicle) and uninfected (Mock/vehicle) samples. Infected ^AX^iAEC treated with remdesivir (SARS-CoV-2/remdesivir) group in the middle. n = 9 – 12 well replicates from 4 chips run in N = 4 independent experiments. **(B)** Pathway enrichment analysis in SARS-CoV-2 infected, vehicle-treated samples compared to Mock/vehicle control reveals JAK-STAT, NF-κB and TNFα as the most upregulated pathways and estrogen, WNT and hypoxia pathway as the most downregulated pathways. n = 10 – 12 well replicates from 4 chips run in N = 4 independent experiments. **(C)** Top upregulated genes in the JAK-STAT pathway upon SARS-CoV-2 infection compared to Mock/vehicle control. n = 10 – 12 well replicates from 4 chips run in N = 4 independent experiments. **(D)** Differentially expressed genes (DEGs) upon SARS-CoV-2 infection compared to mock/vehicle control highlights viral genes, antiviral genes and stress response genes as most significantly upregulated genes in infected wells. n = 10 – 12 well replicates from 4 chips run in N = 4 independent experiments. **(E)** DEGs upon remdesivir treatment in SARS-CoV-2 infected wells compared to Mock/vehicle control highlights viral genes and antiviral genes as most significantly upregulated genes in infected wells. n = 9 – 10 well replicates from 4 chips run in N = 4 independent experiments. **(F)** Number of DEGs in the SARS-CoV-2/vehicle and SARS-CoV-2/remdesivir group compared to Mock/vehicle. n = 9 – 12 well replicates from 4 chips run in N = 4 independent experiments. SCoV-2 = SARS-CoV-2.

Next, we analysed the transcriptional changes that were induced by SARS-CoV-2 infection in ^AX^iAEC and compared the transcriptional profile of infected and mock control groups in the absence of remdesivir treatment. We found strongest upregulation of immune-related and inflammatory pathways (JAK-STAT, NF-κB and TNF-α) and simultaneous downregulation of estrogen, Wnt and hypoxia signalling pathways (**Figure 5B**). This suggests a general impairment of epithelial homeostasis and repair, as indicated by downregulation of Wnt signalling^68,69^ and activation of a pro-inflammatory immune response in infected ^AX^iAEC. Notably, the top upregulated genes in the JAK-STAT signalling pathway were exclusively interferon-stimulated genes (ISGs) with direct antiviral function attributed to IFIT1, IFIT3, RSAD2 and MX1 (**Figure 5C**)^70–73^. Comparing the global transcriptional profile of infected ^AX^iAEC to mock control, we found that pro-inflammatory transcripts were among the top 10 differentially expressed genes (**Figure 5D**). DHX58 and TRIM31 are implicated in RIG-I-like receptor (RLR) signalling and downstream activation of mitochondrial antiviral-signalling protein (MAVS)^74,75^, which results in the activation and nuclear translocation of transcription factors such as IRF3, IRF7 and NF-κB to induce transcription of proinflammatory cytokines and IFNs^76^. The IFN-stimulated genes TRIM31 and PHF11 amplify the immune response^75,77,78^, while EIF2AK2 (also known as proteinkinase R; PKR) inhibits protein translation in infected cells and orchestrates inflammation in response to stress and infection^79^. Moreover, TGM2 and SLC25A28 are implicated in inflammatory responses that are marked by cellular stress, tissue damage and cell death ^80,81^. Little is known about LIM-domain only protein 2 (LMO2) function in alveolar epithelial cells. However, by functional analogy to LIM-domain only protein 4 (LMO4) and its role in tissue repair ^82^, LMO2 might be indicative of an active tissue damage response in the LOC infection model. Notably, LMO2, TGM2 and CEACAM1 regulate immune cell activation^81,83,84^, implying that infected ^AX^iAEC might initiate transcriptional programs to orchestrate cytotoxic cell-mediated immune responses. Interestingly, we detected high expression of CEACAM-1, which mediates immunosuppressive signalling^83^. Finally, we found that the strongest upregulated genes in infected ^AX^iAEC were two viral genes, N and ORF10, implying active viral replication in infected ^AX^iAEC. It is interesting to note, that the viral N and ORF10 genes remain highly expressed in ^AX^iAEC after remdesivir treatment (**Figure 5E**), even though SARS-CoV-2 NP was nearly undetectable by IF staining (**Figure 4B**). However, the host response changed upon remdesivir treatment compared to infected ^AX^iAEC treated with vehicle control. Despite ENSG00000237807 encoding a long non-coding RNA, all other highly upregulated genes in infected, remdesivir-treated ^AX^iAEC encode IFN-stimulated genes (ISG). DHX58 was the only transcript that was also among the top 10 upregulated genes in infected, vehicle-treated ^AX^iAEC and acts as sensor for viral dsRNA to initiate antiviral signalling^74^. In addition, remdesivir treated ^AX^iAEC highly expressed genes that code for proteins with direct antiviral function including MX1, MX2, OAS2 and OASL^73,85–88^. Finally, ^AX^iAEC upregulated ISG15 and ETV7 expression upon remdesivir treatment, which fine-tune IFN response in epithelial cells^89,90^. Taken together, we noted little overlap of differentially expressed genes in infected ^AX^iAEC that were treated with vehicle or remdesivir. A separation of the transcriptional profile also became evident, when analysing for unique and shared differentially expressed genes in ^AX^iAEC that were infected and treated with vehicle control or remdesivir in comparison to mock controls. No transcriptional changes were induced to the host cells by remdesivir treatment itself (0 DEGs specific to SARS-CoV-2/remdesivir group) and only 43 genes remained differentially expressed in infected ^AX^iAEC treated with remdesivir and overlapped with infected ^AX^iAEC treated with vehicle control. In contrast, 1089 genes were dysregulated in the SARS-CoV-2/vehicle group supporting the notion that remdesivir treatment partially reverts SARS-CoV-2-induced transcriptional changes in infected ^AX^iAEC (**Figure 5F**).

In this proof-on-concept study we demonstrated the application of our newly developed LOC SARS-CoV-2 infection model in physiological stretch for drug testing. The antiviral effect of remdesivir was recapitulated in infected ^AX^iAEC/hLMVEC co-cultures protecting the epithelial/endothelial barrier from alveolar damage. Moreover, in-depth molecular characterisation of the model elucidated the antiviral response of ^AX^iAEC upon viral infection and revealed a predominant pro-inflammatory immune and stress response. These transcriptomic alterations were modulated by remdesivir treatment, shifting the transcriptomic profile of infected ^AX^iAEC towards the transcriptional state of uninfected control cells.

## Discussion

In this study, we employed a well-characterised immortalized alveolar epithelial cell line, ^AX^iAEC, to recreate the alveolar epithelium on-chip. We noted a striking impact of ALI on the susceptibility of ^AX^iAEC to SARS-CoV-2 infection. A phenomenon that has already been reported for A549 lung cells^91^ and that we also observed in primary alveolar epithelial cells (^AX^hAEpC). SARS-CoV-2 infection was abortive in submerged ^AX^hAEpC cultures, while low viral titers were released from infected ALI cultures of donor 4. These results confirm our observations in ^AX^iAEC and highlight the importance of alveolar epithelial cell culture in ALI to capture natural susceptibilities to respiratory viruses *in vitro*.

^AX^iAEC ALI cultures were susceptible to SARS-CoV-2 infection, marked by the release of infectious viral progeny and a significant decrease of TER indicative of impaired barrier function at 48hpi. However, following the peak of infection, we noted a decrease in virus production and increase in TER values at 72hpi suggesting clearance of the virus and recovery of barrier function. A similar SARS-CoV-2 replication kinetics in the distal lung was reported in *in vitro* studies using precision-cut lung slices (PCLS)^46,47^ but further investigations are required to better understand the underlying molecular mechanisms of viral clearance. However, we noted a strong upregulation of antiviral genes in ^AX^iAEC upon SARS-CoV-2 infection at 48hpi, particularly of IFNL1 (>5000-fold) and MX1 (>60-fold). It has been shown that SARS-CoV-2 is highly susceptible to IFNs^92,93^, which might explain the clearance of the virus from our model system. Notably, these infection dynamics reflect the clinical situation, which is marked by active viral replication during the acute phase of COVID-19, while later stages of diffuse alveolar damage do not necessarily coincide with detectable SARS-CoV-2 infection and are more likely driven by aberrant activation of immune responses^94–96^. In contrast, ^AX^iAEC monocultures in ALI seem to tightly regulate their immune response to SARS-CoV-2 and downregulated type I and type III IFN expression at 72hpi, which coincided with the release of lower viral titers and recovery from virus-mediated damage. A similar pattern of IFN secretion is thought to be essential for resolution of COVID-19 in mildly and moderately ill patients^76^. Therefore, we hypothesized that the SARS-CoV-2 infection model in ^AX^iAEC monocultures represents characteristics of mild and moderate COVID-19 and only a more physiological model might be able to capture a broader spectrum of COVID-19 manifestations *in vitro*. Therefore, we next employed the ^AX^iAEC/hLMVEC co-culture model in ALI on the LOC. The flexible membrane allows the implementation of cyclic stretch to simulate breathing motion^49^. The application of stretch resulted in barrier formation and TER values in the physiological range^97–99^. We differentiated the ^AX^iAEC/hLMVEC co-culture in ALI and infected them under dynamic or static conditions to evaluate the impact of stretch on SARS-CoV-2 infection in the alveoli. Our results demonstrated that SARS-CoV-2 infection was slightly but significantly more productive in dynamic conditions at 48hpi. Remarkably, SARS-CoV-2-mediated damage was more pronounced under stretch and complete barrier breakdown was achieved in infected cultures at 48hpi. Extended damage to the air-blood barrier upon SARS-CoV-2 infection has also been reported in other LOC models^34,39^. Moreover, we found that stretch is a major factor to modulate the ^AX^iAEC transcriptional profile in response to SARS-CoV-2 infection. Evidence from literature supports our observation that breathing motion modulates the cellular response to infection. Cyclic stretch has been shown to alter cytokine secretion of alveolar epithelial cells^65^ and to shape inflammation in response to nanoparticles, bacteria and influenza virus^63,100^ highlighting the importance of mechanical stimuli to accurately simulate mucosal immunity in the distal lung.

Few studies have addressed SARS-CoV-2 infection of the alveoli on a LOC system. An early study reporting SARS-CoV-2 infection on an alveolus-on-chip employed primary alveolar epithelial cells in co-culture with human lung microvascular endothelial cells incorporating mechanical stimulation (medium flow in the vascular channel and linear stretch of the cell culture membrane). Thacker and colleagues reported unproductive infection of the alveolar system but massive inflammation and endothelial cell activation upon infection. Finally, they identified IL-6 secretion as a driver of the deleterious inflammatory response and suggested tocilizumab, a neutralizing antibody to IL-6 receptor (IL-6R), as a potential drug candidate to reduce damage to the alveolar barrier^39^. This study has provided valuable pre-clinical data contributing to the rapid repurposing and approval of tocilizumab for clinical use in COVID-19 patients. Moreover, it provided encouraging results demonstrating the application of LOC models for pre-clinical drug testing. Other research groups have employed LOC models of the alveoli and airways to test anti-viral compounds blocking SARS-CoV-2 entry or replication. However, technical requirements and sophisticated handling of such complex LOC models hinder their application in BSL3 laboratories. Therefore, most of these studies were conducted with SRAS-CoV-2 pseudovirus, which does not fully recapitulate the entire life cycle of SARS-CoV-2^33,37,101^. In a study using the native virus, Zhang and colleagues infected an alveolus-on-chip model under constant flow with SARS-CoV-2. Infection with high titers of SARS-CoV-2 induced an anti-viral type I IFN response in alveolar epithelial cells and endothelial cell activation. However, only in the presence of peripheral blood mononuclear cells (PBMC), SARS-CoV-2 infection resulted in barrier disruption and leakage^43^. In contrast, SARS-CoV-2 infection critically affected barrier tightness at lower MOIs and even in the absence of immune cells in our newly developed LOC model incorporating physiological stretch.

Transcriptomic analysis of infected ^AX^iAEC in the breathing LOC infection model revealed the activation of antiviral and inflammatory signalling pathways upon SARS-CoV-2 infection. The downregulation of estrogen signalling pathway may further contribute to an inflammatory response in the infected LOC system. Estrogen signalling mediates anti-inflammatory responses in viral infections, modulates the renin-angiotensin system^102^ and is directly modulated by SARS-CoV-2 Spike (S) protein^103^. Notably, DHX58 is consistently upregulated in infected ^AX^iAEC and, together with MDA5, is the main receptor to sense SARS-CoV-2 RNA in lung epithelial cells ^104^ resulting in NF-κB activation and subsequent expression of type I and III interferon-stimulated genes (ISGs) such as TRIM31 and PHF11^77,78,105^. Multiple of the top differentially expressed genes (DEGs) in our LOC model are implicated in COVID-19 and are either dysregulated upon SARS-CoV-2 infection or targeted by the virus to interfere with the host antiviral immune response. For example, the interaction of TRIM31 with MAVS is targeted by SARS-CoV-2 to attenuate the host’s antiviral defence mechanisms^106^. Similarly, EIF2AK2 has been consistently found to be upregulated during SARS-CoV-2 infection and was identified as a direct target of the viral N protein^107^. Furthermore, increased expression of the immune-suppressive receptor CEACAM-1 has been detected in nasal swabs, autopsies and blood of COVID-19 patients and correlated with disease severity^108^. Whether upregulation of TGM2 is associated with severe COVID-19 remains controversial^109,110^. However, Occhigrassi and colleagues demonstrated that TGM2 negatively regulated STING pathway and reduced production of IFN-β, which might contribute to the delayed IFN response in COVID-19 patients^110^. An early anti-viral IFN response is crucial to fight SARS-CoV-2 infection and a dysfunctional IFN response due to anti-IFN autoantibodies or genetic predispositions has been linked to critical COVID-19^111,112^. However, elevated IFN levels at advanced stages of COVID-19 may contribute to moderate or severe disease outcome^113,114^. Whether IFN production is impaired in the infected ^AX^iAEC, and whether the LOC model would progress to immune misfiring resembling the cytokine storm in severe COVID-19 patients requires further investigations. We hypothesize that our LOC infection model captures the early phase of SARS-CoV-2 infection that is marked by inflammation and cellular stress response. The early stages of severe COVID-19 are an interesting therapeutic window for the development of targeted drugs to guide medical intervention in high-risk patients before they progress to life-threatening disease stage. Potentially, the LOC infection model might serve as a valuable *in vitro* tool for pre-clinical drug testing for COVID-19 and other respiratory virus disease. Therefore, we developed a drug treatment protocol on the LOC system using the FDA-approved antiviral drug remdesivir. It was the first effective antiviral treatment for hospitalized COVID-19 patients^115^ and its activity has been demonstrated in microphysiological systems (MPS) of SARS-CoV-2 infection in the upper^44^ and lower respiratory tract^43,45^. In line with previous reports, remdesivir treatment in our medium-throughput ^AX^iAEC model on cell culture inserts was highly effective and fully protecting the alveolar barrier from SARS-CoV-2-mediated damage at doses in the micromolar range. Similarly, 1μM remdesivir treatment inhibited SARS-CoV-2 replication in the infected LOC system and protected the alveolar barrier from excessive damage upon infection. In line with our results, Zhang and colleagues showed that remdesivir treatment interfered with SARS-CoV-2 replication and partially restored the alveolar epithelial/endothelial barrier in an alveolus-on-chip model even in the presence of PBMC that contribute to exacerbate inflammation^43^. Notably, transcriptomic analysis of infected ^AX^iAEC demonstrated a partial reversal of the inflammatory response towards reconstitution of the uninfected state upon remdesivir treatment in our model system. The top 10 DEGs in remdesivir treated ^AX^iAEC are nearly all ISGs with well described functions for OASL, OAS2, MX1 and MX2 in the inhibition of viral replication and degradation of viral RNA ^73,85–90^. This suggests that remdesivir treatment induces a robust IFN response in the infected LOC. Notably, remdesivir treatment reduced blood cytokine levels in COVID-19 patients^116,117^ but little is known about its impact on alveolar immune responses due to limitations in sample acquisition. The human-derived LOC model could overcome these limitations allowing to study underlying molecular mechanisms at the primary site of drug action and contribute to improved patient stratification and therapy selection. Furthermore, organ-on-chip technology holds the potential to revolutionize drug development enabling drug testing in a human-relevant context to develop safer and more effective drugs while at the same time reducing costs and accelerating drug development. Particularly in light of global efforts to reduce animal experimentation, the development of MPS of different organs and diseases as a part of New Approach Methodologies (NAMs) increasingly gains importance. Recently, Hesperos Inc. (Orlando, USA) has leveraged human-on-a-chip technology to test efficacy of a new monoclonal antibody against rare neuropathies, a disease that cannot be modelled in experimental animals, and demonstrated a therapeutic benefit leading to FDA-approval for the new therapy ^118^. Another milestone in the transition from animal models to human-derived models in drug development has been achieved with the latest FDA modernization ACT 3.0. In April 2025, the FDA announced to phase out the requirement for animal testing for the development of monoclonal antibodies and selected drugs^119^. This new regulation might also accelerate drug development for COVID-19 with two monoclonal antibodies (Tocilizumab and Vilobelimab) already being approved for clinical use in COVID-19 patients. In future, safety studies in animal models might become dispensable, and LOC technology may serve as a useful tool for safety and efficacy testing in the development of novel antibody therapies for COVID-19 patients.

Nevertheless, our LOC infection model meets some important limitations. Firstly, we have focused our investigations on the acute phase of SARS-CoV-2 infection but did not employ infected chips for long-term experiments to study prolonged disease courses and alveolar recovery after viral clearance. Second, we have not incorporated immune cells into the system despite alveolar macrophages and PBMCs playing an essential role in the pathogenesis of severe COVID-19^120^. Particularly considering our transcriptomic data demonstrating that ^AX^iAEC upregulate genes that regulate cytotoxic immune cells, incorporation of these cell types into the LOC could further exacerbate lung injury and contribute to the biological relevance of our infection model. Finally, we established the LOC infection model using cell lines. Despite closely resembling primary human alveolar epithelial cells, the immortalization process might affect behaviour of ^AX^iAEC in unknown but biologically relevant manners. Moreover, genetic factors and pre-existing comorbidities among patients affect the risk to develop severe COVID-19^121^. Such interpatient differences are not considered in our model system because we decided to resign from using primary alveolar epithelial cells in the LOC due to their heterogeneous behaviour *in vitro*. The application of the ^AX^iAEC cell line allowed us to generate a reproducible and easy-to-handle system, which facilitates application of our newly developed SARS-CoV-2 infection LOC model for lung research, virology, and pre-clinical safety and efficacy testing.

## Conclusion

Taken together, we demonstrated the importance of ALI to recapitulate natural susceptibilities of alveolar epithelial cells to respiratory viruses *in vitro* and developed a SARS-CoV-2 infection model on a LOC device under physiological stretch. This model mimicking breathing motion supported SARS-CoV-2 infection and recapitulated key aspects of acute COVID-19, such as lung damage and inflammation in response to the intrinsic defence mechanisms of alveolar epithelial cells to viral infection. Finally, in this proof-of-concept study, we have used the LOC infection model for drug testing to expand the toolbox of human-relevant alveolar models for pre-clinical applications. This may accelerate pre-clinical drug development for respiratory infectious diseases and elevate global preparedness to face novel emerging virus variants.

## Acknowledgements

We would like to thank Dr. Kathrin Summermatter, Monika Gsell-Albert, and Julia Feldmann from the Institute for Infectious Diseases (IFIK), University of Bern, for their support involving any work in the BSL3 facility, protocol approval and implementation of new methods that guaranteed rapid transfer and operation of the LOC system in a BSL3 setting.

## Author contribution

Conceptualization: MK, NR, NH, MKJ, TG. Experiment design: MK, NR, LdM, NH, MKJ, MDM. Experiment conduction: MK. Data analysis: MK, PC. Resources: LdM, LF, JS. Writing – original draft: MK. Writing – revision & editing: MK, LdM, NR, PC, NH, MKJ. All authors read and approved the final manuscript. Supervision: NH, MKJ. Project Administration: MK, LdM, NR, LF, JS, NH, MKJ. Funding Acquisition: TG, MKJ, NH, MK.

