## Supplemental Material for "A Human Alveolus-on-Chip Recapitulates SARS-CoV-2-mediated Lung Injury in an Organ-relevant Context for Pre-clinical Applications"

### Supplementary Figure 1

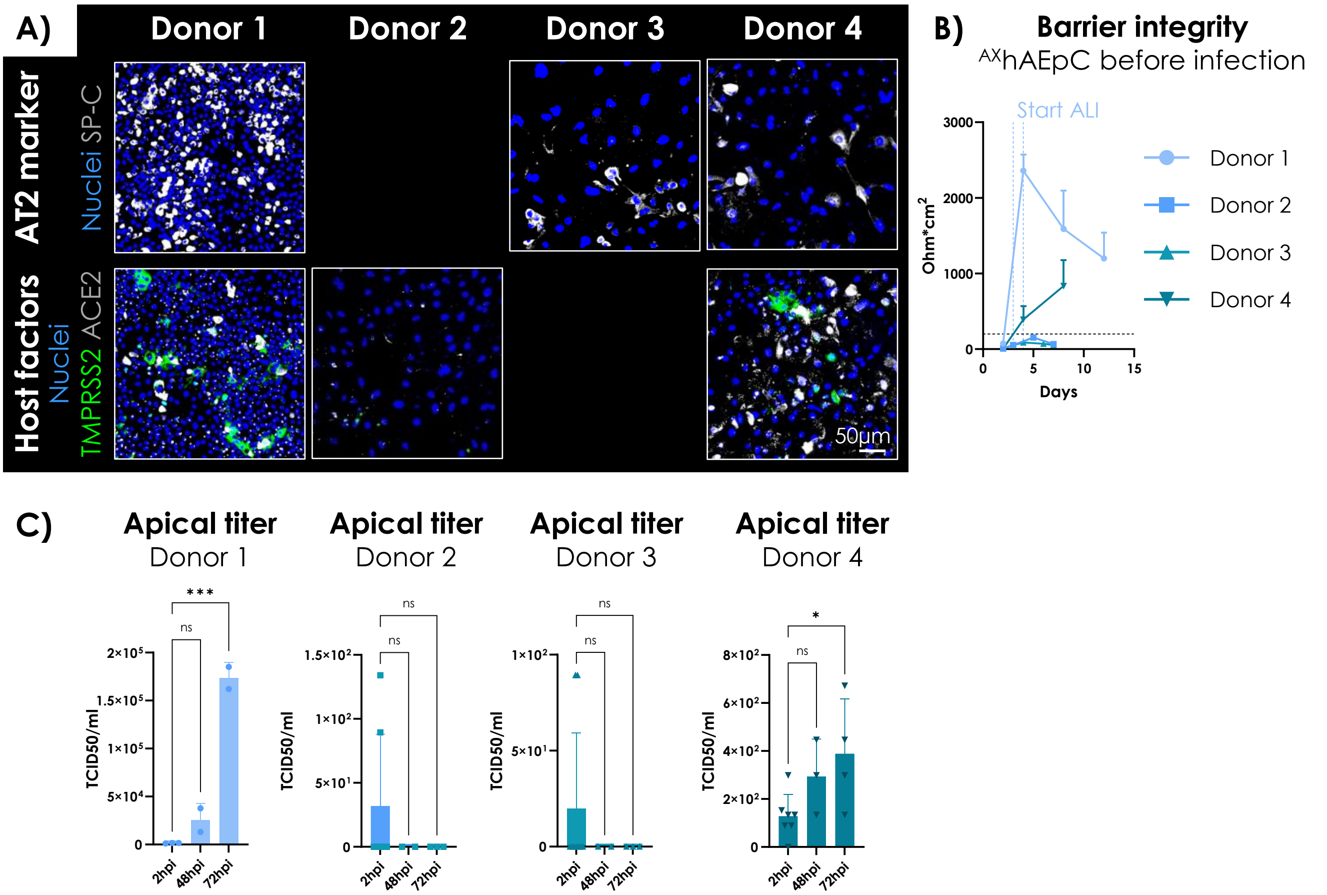


**Supplementary Figure 1:** ***SARS-CoV-2 infection dynamics in primary $^{AX}$hAEpC are donor-dependent.***
**(A)** Primary $^{AX}$hAEpC were isolated from 4 donors and seeded on cell culture inserts for maturation in ALI culture. Samples were fixed for IF staining between day 8 and day 15. Expression of AT2 marker SP-C (grey) and host factors ACE2 (grey) and TMPRSS2 (green) was evaluated. Nuclei were stained with Hoechst-33342 (blue). Representative images are shown. Not all markers could be evaluated for donor 2 and donor 3. Scale bar = 50µm. **(B)** $^{AX}$hAEpC were differentiated in ALI culture starting between day 3 – 4 in culture (dotted lines, blue) and TER was measured every 2 – 4 days. A threshold of 200 Ohm*cm$^2$ (dotted line, black) was defined as tight alveolar barrier. n = 6 – 23 replicates per donor. **(C)** SARS-CoV-2 titer in ALI cultures of $^{AX}$hAEpCs was determined from apical washes at 2hpi, 48hpi and 72hpi for each donor. n = 2 – 9 replicates per donor. Statistics: Ordinary one-way ANOVA with Dunnett's multiple comparison (with a single pooled variance) post hoc test. *, $p \leq 0.05$; ***, $p \leq 0.001$.

### Supplementary Figure 2

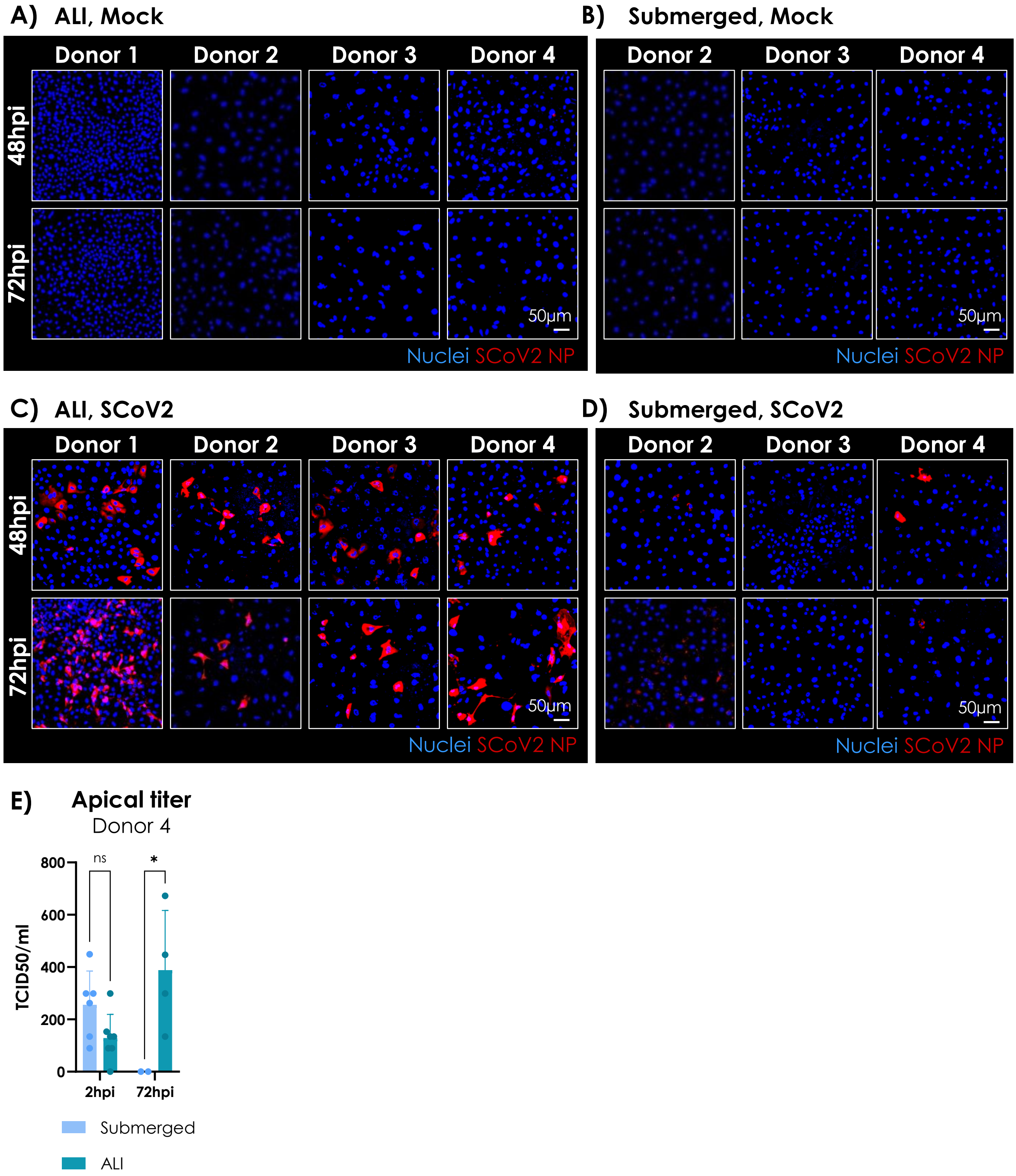


**Supplementary Figure 2:** ***ALI culture supports SARS-CoV-2 infection in $^{AX}$hAEpC.***
**(A) & (B)** $^{AX}$hAEpC were cultured in ALI (A) or submerged (B) condition and were treated as mock control. Cells were fixed at 48hpi or 72hpi and stained for SARS-CoV-2 (SCoV2) NP (red). Nuclei were counterstained with Hoechst-33342 (blue). Representative images of each tested donor are shown. Scale bar = 50µm. **(C) & (D)** $^{AX}$hAEpC were cultured in ALI (C) or submerged (D) condition and infected with SARS-CoV-2 at MOI 0.5. Cells were fixed at 48hpi or 72hpi and stained for SARS-CoV-2 (SCoV2) NP (red). Nuclei were counterstained with Hoechst-33342 (blue). Representative images of each tested donor are shown. Scale bar = 50µm. **(E)** Apical washes were collected from $^{AX}$hAEpC of donor 4 at 2hpi and 72hpi to compare SARS-CoV-2 production under submerged and ALI condition. Apical SARS-CoV-2 titer was determined on Vero E6 cells and TCID50/ml was calculated. N = 1 donor; n = 2 – 7 replicates. Statistics: Ordinary two-way ANOVA with Sidak's multiple comparison (with a single pooled variance) post hoc test. *, $p \leq 0.05$.

### Supplementary Figure 3

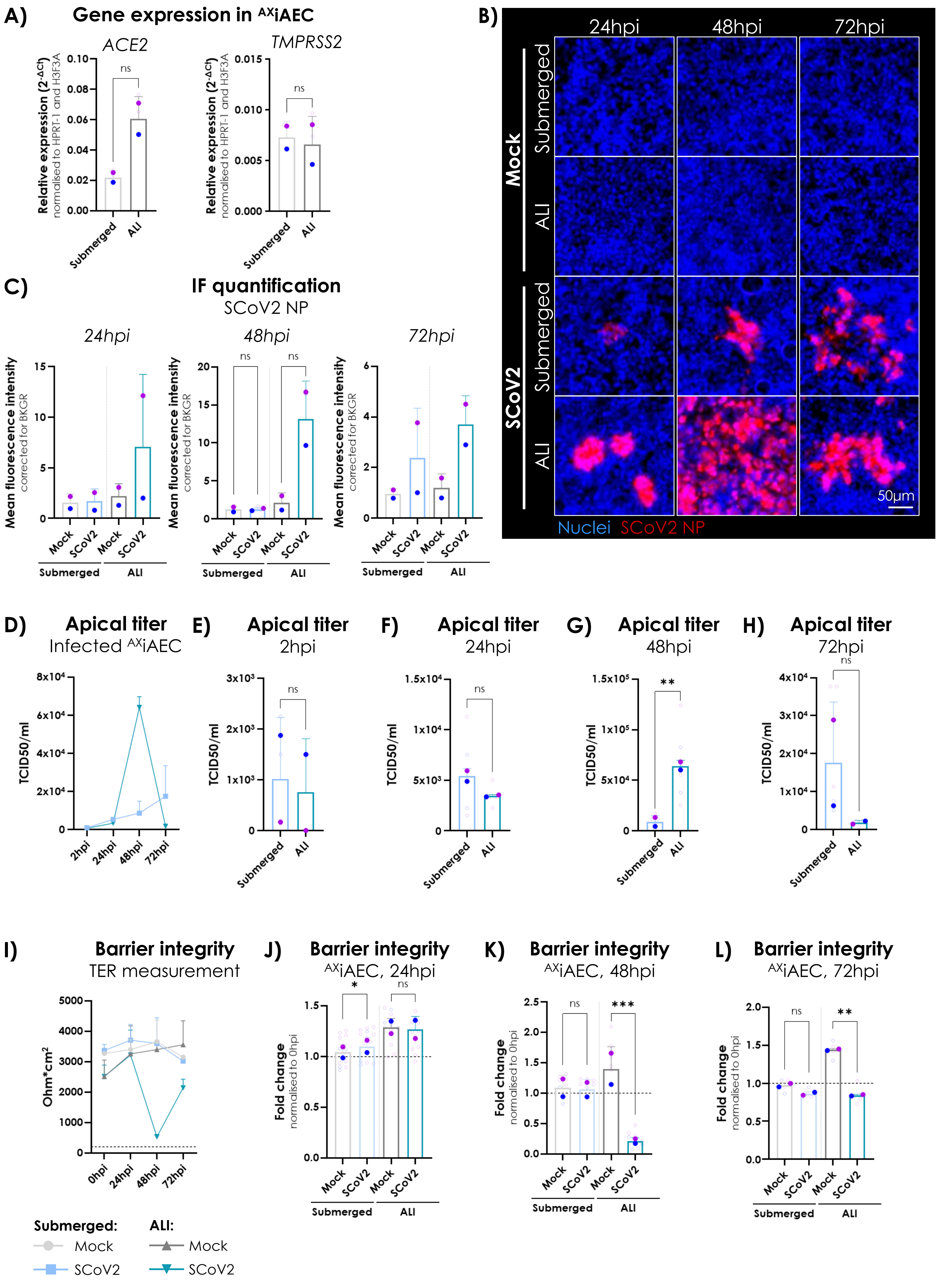

**Supplementary Figure 3:** ***SARS-CoV-2 infection of*** $^{AX}$***iAEC peaks at 48hpi in ALI culture.***
**(A)** $^{AX}$iAEC were differentiated on cell culture inserts in submerged or ALI conditions for more than 28 days before cell lysis and gene expression analysis of ACE2 and TMPRSS2. Relative expression was normalized to housekeeping genes HPRT-1 and H3F3A. n = 3 replicates run in N = 2 independent experiments. Statistics: Paired t-test. ns, $p > 0.05$. **(B)** $^{AX}$iAEC on cell culture inserts were fixed at 24hpi, 48hpi and 72hpi and infection efficacy in submerged and ALI cultures was evaluated by IF staining for SARS-CoV-2 NP (SCoV2 NP; red). Nuclei were stained with Hoechst-33342 (blue). No virus was detected in mock control while SCoV2 NP was detected in infected cultures in submerged and ALI conditions at all sampled time points. Representative images are shown. Scale bar = 50µm. **(C)** Image quantification for corrected MFI of SCoV2 NP staining at 24hpi (left), 48hpi (middle) and 72hpi (right). N = 2 independent experiments; n = 1 – 2 replicates per experiment. Statistics for 48hpi time point: Ratio paired t-test. ns, $p > 0.05$. No statistics performed at 24hpi and 72hpi due to small sample size (n < 3 replicates). **(D)** Apical virus titer was determined at 2hpi, 24hpi, 48hpi and 72hpi in $^{AX}$iAEC differentiated on cell culture inserts in submerged or ALI cultures and infected with SARS-CoV-2 at an MOI 0.5. N = 2 independent experiments; n = 1 – 3 replicates per experiment. **(E) – (H)** SARS-CoV-2 apical titer of infected $^{AX}$iAEC was compared for submerged and ALI cultures at (E) 2hpi, n = 2 – 4 replicates; at (F) 24hpi, n = 4 – 5 replicates; at (G) 48hpi, n = 5 replicates; at (H) n = 3 – 5 replicates. N = 2 independent experiments in all plots. Statistics: Paired t-test. ns, $p > 0.05$; **, $p \leq 0.01$. **(I)** Barrier integrity of mock and SARS-CoV-2 infected $^{AX}$iAEC in submerged and ALI cultures was evaluated by TER measurement at 0hpi, 24hpi, 48hpi and 72hpi. A threshold of TER ≥200 Ohm*cm$^2$ (dotted line) was considered as tight barrier. n = 4 – 14 replicates run in N = 2 experiments. **(J) – (L)** Relative TER values are represented as fold change to 0hpi for (J) 24hpi, n = 11 – 13 replicates; (K) 48hpi, n = 6 – 10 replicates; (L) 72hpi, n = 4 – 5 replicates. N = 2 independent experiments in all plots. Statistics: Ratio paired t-test. ns, $p > 0.05$; *, $p \leq 0.05$; **, $p \leq 0.01$; ***, $p \leq 0.001$.

### Supplementary Figure 4

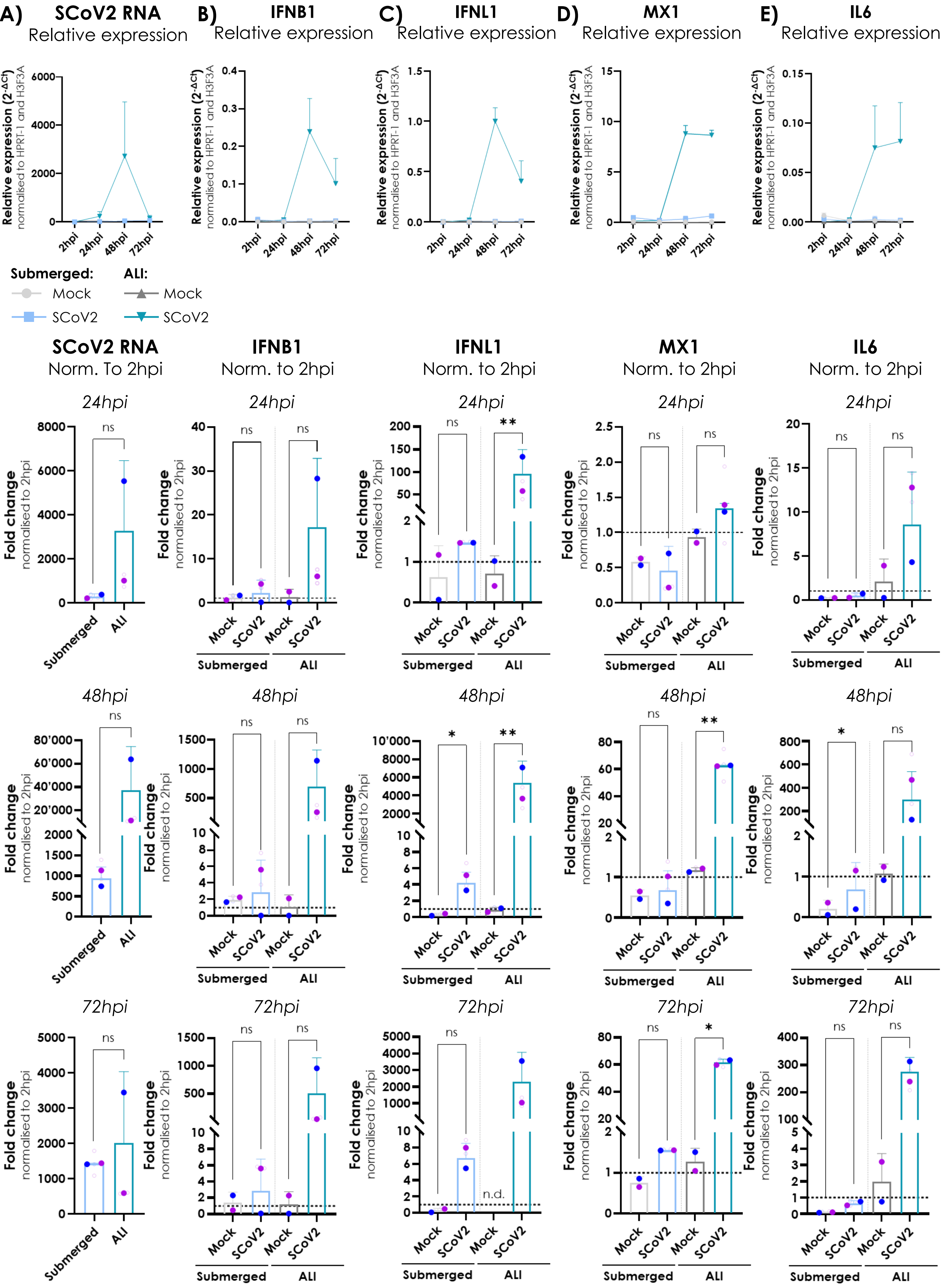

**Supplementary Figure 4:** ***ALI shapes the innate immune response of $^{AX}$iAEC to SARS-CoV-2.***

**(A) – (E)** *Top*: Relative expression of (A) SARS-CoV-2 (SCoV2) viral RNA, (B) IFNB1, (C) IFNL1, (D) MX1 and (E) interleukin-6 (IL6) in infected $^{AX}$iAEC lysates was analyzed at 2hpi, 24hpi, 48hpi and 72hpi. *Bottom panels*: Fold change to 2hpi was calculated for each tested gene at 24hpi, 48hpi and 72hpi. n = 2 – 3 replicates run in N = 2 independent experiments. Statistics: Paired t-test. ns, $p > 0.05$; *, $p \leq 0.05$; **, $p \leq 0.01$. n.d. = not determined. SCoV2 = SARS-CoV-2.

### Supplementary Figure 5

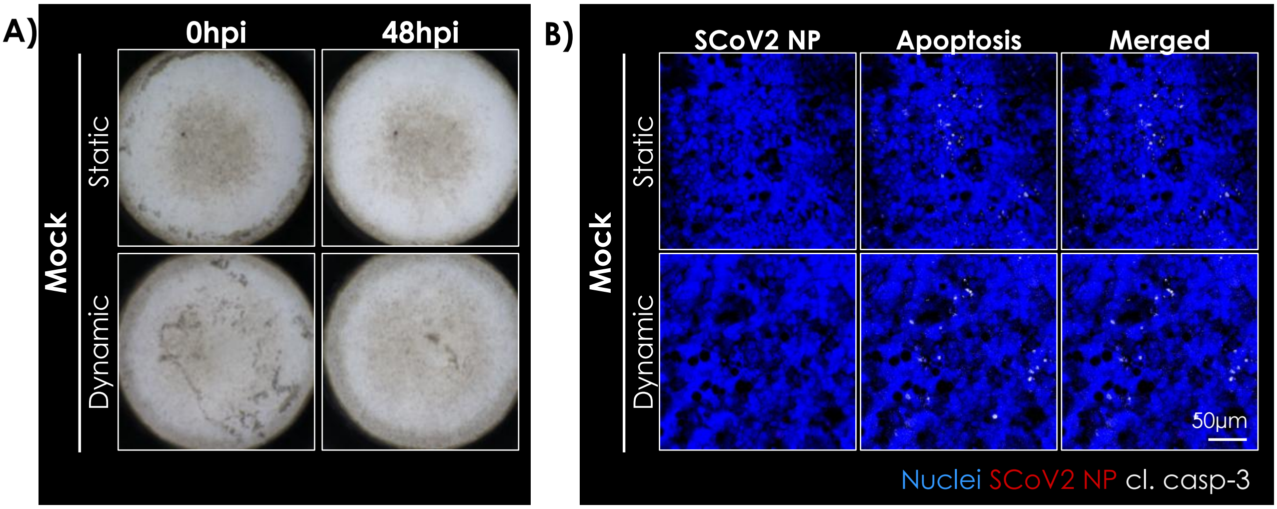


**Supplementary Figure 5:** ***Characterization of the SARS-CoV-2 infection model on the LOC system.***
**(A)** Representative images of mock control wells in ALI/static and ALI/dynamic at 0hpi and 48hpi. **(B)** LOC in ALI/static and ALI/dynamic was fixed and stained for SARS-CoV-2 (SCoV2) NP (red) and cl. casp-3 (grey) at 48hpi. Nuclei were counterstained with Hoechst-33342 (blue). No signal for SCoV2 NP was observed and only few apoptotic cells were found in mock control. Representative images are shown. Scale bar = 50µm.

### Supplementary Figure 6

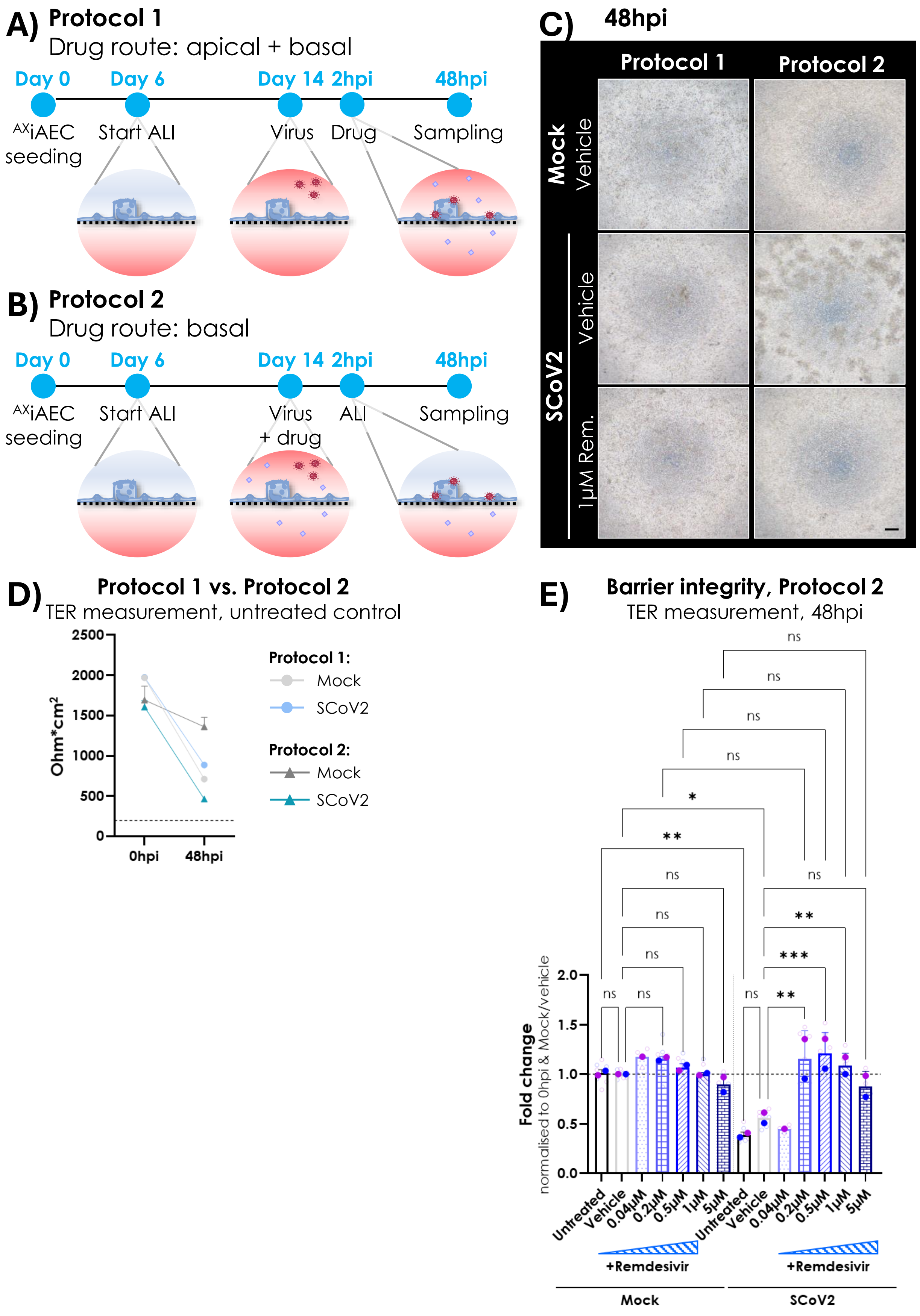


**Supplementary Figure 6:** ***Development of a drug treatment protocol for $^{AX}$iAEC on cell culture inserts.***
**(A)** Schematic illustration of infection and drug treatment protocol 1: $^{AX}$iAEC were differentiated on cell culture inserts in ALI starting on day 6. $^{AX}$iAEC were infected on day 14 with SARS-CoV-2 at MOI 0.5. The inoculum was removed after 2h incubation and treatment with remdesivir (0.2µM, 0.5µM, 1µM and 5µM) or vehicle control (0.5% DMSO) started from the apical and basal side. During this period, cells were cultured in submerged conditions for 48h until samples were harvested at experiment endpoint.

**Supplementary Figure 6:** ***continued***

**(B)** Schematic illustration of infection and drug treatment protocol 2: $^{AX}$iAEC were differentiated on cell culture inserts in ALI starting on day 6. $^{AX}$iAEC were infected on day 14 with SARS-CoV-2 at MOI 0.5. At the same time, remdesivir (0.04µM, 0.2µM, 0.5µM, 1µM and 5µM) or vehicle control (0.5% DMSO) was administered from the apical and basal side. The inoculum was removed after 2h incubation and ALI culture was restored. Treatment was continued from the basal side for 48h, subsequently, samples were harvested. **(C)** $^{AX}$iAEC monocultures were infected and treated on cell culture inserts according to protocol 1 or 2, respectively, and were evaluated for cytopathic effect (CPE) at 48hpi. Representative bright field images are shown. Scale bar = 200µm. **(D)** Barrier integrity of untreated $^{AX}$iAEC infected according to protocol 1 or protocol 2 was assessed by TER measurement at 0hpi and 48hpi. Employing protocol 1, TER values decreased from approx. 2000 Ohm*cm$^2$ to < 1000 Ohm*cm$^2$ in SARS-CoV-2 infected and mock control inserts. Instead, TER in mock infected $^{AX}$iAEC remained stable > 1000 Ohm*cm$^2$ at 48hpi and only dropped in SARS-CoV-2 infected inserts to TER values < 500 Ohm*cm$^2$ when employing protocol 2. The barrier remained intact in all conditions with TER values > 200 Ohm*cm$^2$ (dotted line, black). N = 2 independent experiments for protocol 2 and N = 1 experiment for protocol 1; n = 4 replicates per experiment. **(E)** Relative TER values at 48hpi are represented as fold change to 0hpi and Mock/vehicle control for $^{AX}$iAEC infected and treated according to protocol 2. The impact of SARS-CoV-2 infection on barrier integrity with or without remdesivir treatment was evaluated. N = 1 – 2 independent experiments; n = 2 – 4 replicates per experiment. Statistics: Repeated-measures one-way ANOVA without Geisser-Greenhouse correction with Sidak's multiple comparison (with a single pooled variance) post hoc test. ns, $p > 0.05$; *, $p \leq 0.05$; **, $p \leq 0.01$; ***, $p \leq 0.001$. SCoV2 = SARS-CoV-2.

### Supplementary Figure 7

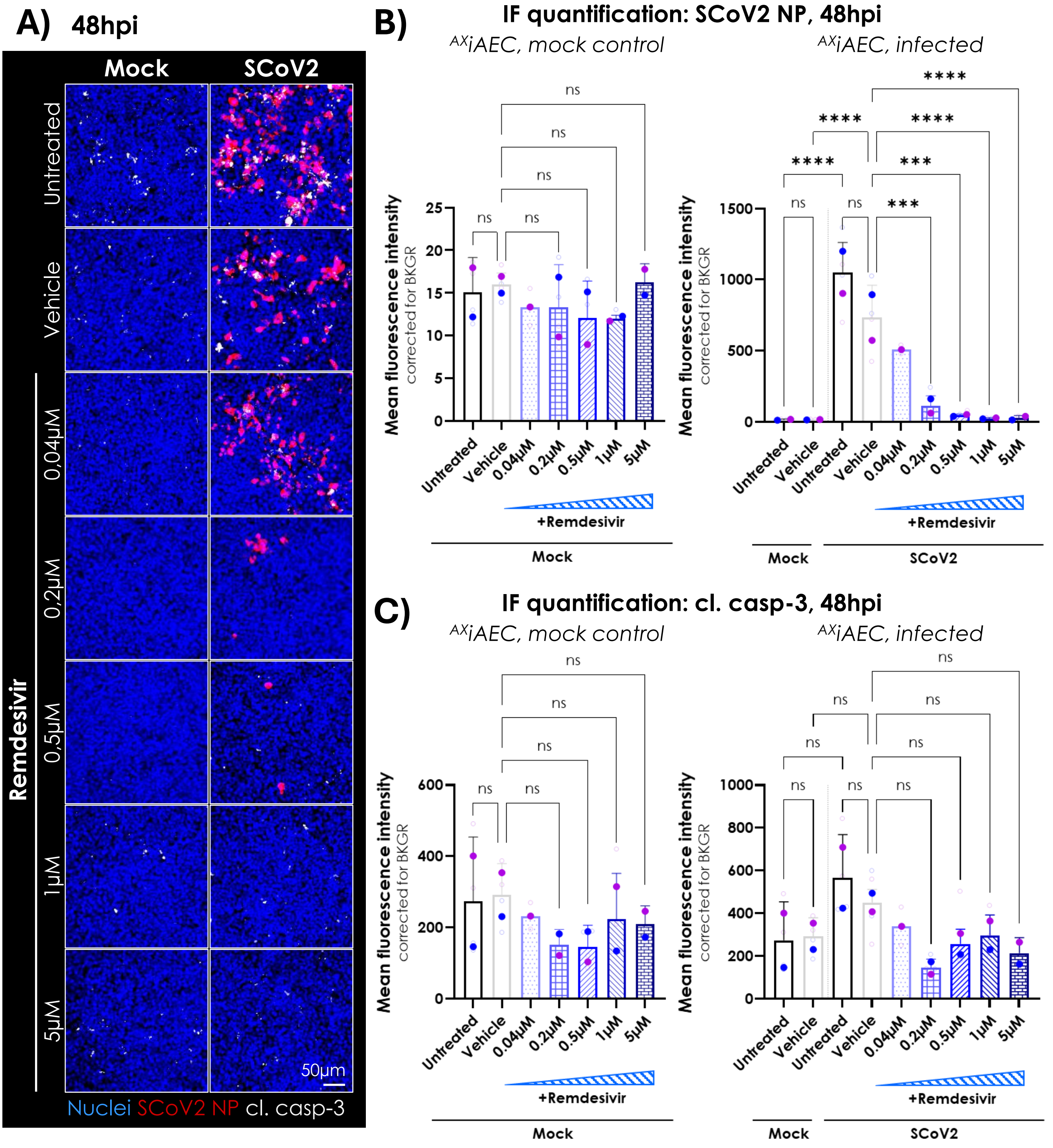


**Supplementary Figure 7:** ***Low doses of remdesivir reduce viral load in AXiAEC ALI cultures on inserts.***
**(A)** AXiAEC infected and treated according to protocol 2 were fixed at 48hpi. SARS-CoV-2 infection efficacy and apoptotic cell death was evaluated by IF staining for SARS-CoV-2 (SCoV2) NP (red) and cl. casp-3 (grey) in mock control and SARS-CoV-2 infected inserts that were untreated or treated with vehicle control (0.5% DMSO) or remdesivir (0.04µM, 0.2µM, 0.5µM, 1µM and 5µM). Representative images are shown. Scale bar = 50µm. **(B)** Quantification of corrected mean fluorescence intensity (MFI) for SCoV2 NP in mock control (left) and infected AXiAEC (right) at 48hpi. Signal intensities were compared to respective vehicle control. N = 1 – 2 independent experiments; n = 1 – 2 replicates per experiment. Statistics: Repeated-measures one-way ANOVA without Geisser-Greenhouse correction with Sidak's multiple comparison (with a single pooled variance) post hoc test. ns, $p > 0.05$; ***, $p \leq 0.001$; ****, $p \leq 0.0001$. **(C)** Quantification of corrected mean fluorescence intensity (MFI) for cl. casp-3 in mock control (left) and infected AXiAEC (right) at 48hpi. Signal intensities were compared to respective vehicle control. N = 1 – 2 independent experiments; n = 1 – 2 replicates per experiment. Statistics: Repeated-measures one-way ANOVA without Geisser-Greenhouse correction with Sidak's multiple comparison (with a single pooled variance) post hoc test. ns, $p > 0.05$. BKGR = background; SCoV2 = SARS-CoV-2.

***Supplementary Table 1:*** *Antibodies.*

| Antibody | Source | Identifier | Dilution |
|---|---|---|---|
| **Mouse anti-SARS-CoV-2 nucleocapsid protein (SCoV2 NP)** | Sino Biologicals | Cat#40143-MM05-100 | 1:1000 |
| **Rabbit anti-cleaved caspase-3 (cl. casp-3)** | Cell Signaling | Cat#9661 | 1:500 |
| **Rabbit anti-mature SP-C** | Seven Hills | Cat#WRAB-76694 | 1:100 |
| **Mouse anti-HTI-56** | Terrace Biotech | Cat# TB-29 AHT1-56 | 1:50 |
| **Rabbit anti-VE-cadherin** | Invitrogen | Cat#36-1900 | 1:100 |
| **Rabbit anti-ACE2** | Abcam | Cat#ab15348 | 1:500 |
| **Mouse anti-TMPRSS2-AF488** | Santa Cruz | Cat# sc-515727 AF488 | 1:50 |
| **Mouse anti-ZO-1-AF488** | Invitrogen | Cat#33-9188 | 1:100 |
| **AlexaFluor 555 donkey anti-mouse IgG** | Invitrogen | Cat#A31570 | 1:200 |
| **AlexaFluor 647 donkey anti-rabbit IgG** | Invitrogen | Cat# A31573 | 1:200 |

***Supplementary Table 2:*** *Primer sequences.*

| Target | Forward primer | Reverse primer | Reference |
|---|---|---|---|
| **IFNB1** | ATGACCAACAAGTGTC TCCTCC | GGAATCCAAGCAA GTTGTAGCTC | PrimerBank ID: 50593016c1 |
| **IFNL1** | GGTGACTTTGGTGCTA GGCT | TGAGTGACTCTTC CAAGGCG | In-house designed (PrimerBlast) |
| **MX1** | GGTGGTCCCCAGTAAT GTGG | CGTCAAGATTCCG ATGGTCCT | PrimerBank ID: 222136618c3 |
| **IL6** | CCTGAACCTTCCAAAG ATGGC | TTCACCAGGCAAG TCTCCTCA | PrimerBank ID: 224831235c2 |
| **SARS-CoV-2 N1** | GAC CCC AAA ATC AGC GAA AT | TCT GGT TAC TGC CAG TTG AAT CTG | Lu et al. 2020[1] |
| **ACE2** | CGAAGCCGAAGACCTG TTCTA | GGGCAAGTGTGGA CTGTTCC | In-house designed (PrimerBlast) |
| **TMPRSS2** | GGACAGTGTGCACCTC AAAGAC | TCCCACGAGGAAG GTCCC | In-house designed (PrimerBlast) |
| **HPRT-1** | AGACTTTGCTTTCCTTG GTCAGG | GTCTGGCTTATATC CAACACTTCG | In-house designed (PrimerBlast) |
| **H3F3A** | GGT GTC TTC AAA AAG GCC AA | GCG AGA AAT TGC TCA GGA CT | In-house designed (PrimerBlast) |

1. Lu X, Wang L, Sakthivel SK, et al. US CDC Real-Time Reverse Transcription PCR Panel for Detection of Severe Acute Respiratory Syndrome Coronavirus 2. *Emerg Infect Dis*. 2020;26(8):1654-1665. doi:10.3201/eid2608.201246

***Supplementary Table 3****: Patient information of $^{AX}$hAEpC donors.*

| | Age | Sex | Commorbidities | Smoker |
|---|---|---|---|---|
| **Donor 1** | 61y | Female | NSCLC, COPD, suspicion of autoimmune disease | Unknown |
| **Donor 2** | 66y | Female | Suspicion for NSCLC | Smoker (40 py) |
| **Donor 3** | 53y | Female | NSCLC | No smoker |
| **Donor 4** | 66y | Male | NSCLC | Former smoker |
